# Antiretroviral treatment does not prevent extrapulmonary tuberculosis during SIV/Mtb co-infection in macaques

**DOI:** 10.1101/2025.08.21.671578

**Authors:** Collin R. Diedrich, Tara Rutledge, Janelle Gleim, Christopher Kline, Pauline Maiello, Jessica Medrano, H. Jacob Borish, Harris Chishti, Justin Gaines, Edwin Klein, Forrest Hopkins, Jacob Klein, Daniel Fillmore, Kara Kracinovsky, Jaime Tomko, Jennifer Schober, Sarah Fortune, Michael Chao, JoAnne L. Flynn, Zandrea Ambrose, Philana Ling Lin

## Abstract

Co-infection with both HIV and *M. tuberculosis* (Mtb) results in disseminated tuberculosis (TB) and accelerated progression of HIV. Despite greater access to antiretroviral treatment (ART), it remains unclear whether suppression of HIV replication protects against severe Mtb infection. Here, using a macaque model of SIV/Mtb coinfection, we investigated whether treatment of SIV infection with ART influenced control of a subsequent Mtb challenge compared to SIV infected macaques who were not treated with ART. Using a macaque model of simian immunodeficiency virus (SIV)-Mtb co-infection, macaques were first infected with SIV_B670_, SIV_B670_ with ART, or saline followed by a low-dose Mtb inoculation with serial clinical, microbiological, PET CT imaging, and immunologic assessments. At necropsy, gross pathology, viremia, bacterial burden, and immunologic parameters were compared. SIV-TB animals had greater gross pathology and total bacterial burden than TB only and SIV/ART/TB groups. However, despite normal blood CD4 counts and undetectable SIV RNA, SIV/ART/TB macaques showed similar clinical parameters and extrapulmonary involvement as SIV/TB animals. Analysis of barcoded-Mtb suggests ART control of SIV replication does not prevent Mtb extrapulmonary dissemination. These data indicate that people living with HIV on ART remain at high risk of bacterial dissemination and extrapulmonary TB disease, particularly when methods to identify extrapulmonary disease are inconsistent. This highlights the importance of understanding the mechanism of extrapulmonary spread and disease severity in HIV/TB co-infected individuals.

## Introduction

Tuberculosis (TB), the disease caused by *Mycobacterium tuberculosis* (Mtb), is currently the most common infectious cause of death, leading to 1.25 million deaths in 2023 with 10.8 million new active TB cases (1). Human immunodeficiency virus (HIV) is the most significant risk factor for TB, even when CD4 T cell counts are in the normal range, and HIV/Mtb co-infection accelerates progression of both TB and AIDS (2). Clinical manifestations of TB among people living with HIV (PLWHIV) often vary by degree of immune suppression. While pulmonary disease remains the most common clinical manifestation of TB, atypical presentations can occur and are often associated with low CD4 counts. Symptoms are often more subtle, chest radiographs can have atypical findings with negative sputum smears, and the rates of extrapulmonary disease (defined as TB pathology at anatomical sites outside of the thoracic cavity) are significantly higher than HIV-naive individuals (3–5). The presence of extrapulmonary TB is more likely to be diagnosed in HIV/Mtb co-infected people and is associated with increased mortality (6).

Antiretroviral treatment (ART) to control HIV replication has dramatically improved HIV and HIV/Mtb co-infection care, especially as it is now available in up to 75% of PLWHIV (7). Delays in ART and low CD4 T cell counts correlate with increased TB incidence over time (8–10) and ART has been shown to reduce mortality from TB and the overall incidence of TB (11–13). Among HIV/Mtb co-infected individuals, ART can prevent or remedy the loss of CD4 T cells (9, 14, 15), improve Mtb-specific CD4 and CD8 T cell responses (16), and reduce macrophage dysfunction (17). However, ART only partially ameliorates the HIV-induced increase in susceptibility to TB (18), as PLWHIV on ART are still more susceptible to TB than HIV-uninfected individuals (10, 19, 20). The mechanisms by which HIV increases susceptibility to TB and why ART only partially reduces that risk are not well understood. It was initially presumed that a low CD4 count was the primary cause of the increase in TB susceptibility in PLWHIV (15), yet we and others have shown that there are CD4-independent risks associated with SIV-induced TB pathogenesis (21–25). Human cohort studies are challenging to study this, as the order in which the infections occur (i.e., primary vs secondary HIV infection) and the duration of each infection are often unknown. For example, acute Mtb infection after chronic HIV infection may lead to worse severity than longstanding asymptomatic Mtb infection (latent) and subsequent acute HIV infection. To dissect these factors, highly controlled animal models to interrogate the phenomena and mechanisms of pathogenesis are beneficial.

Non-human primate (NHP) models of TB recapitulates many key features of human Mtb infection (26, 27) and we and others have shown that they can be productively infected with SIV to better understand HIV-Mtb co-infection (21, 22, 24, 25, 28, 29). Here, cynomolgus macaques were infected with SIV with or without ART and subsequently infected with Mtb to determine the influence of SIV replication or ART suppression of SIV on susceptibility to acute Mtb infection, using a barcoded strain of virulent Mtb. Our data confirm that SIV increases TB pathology and influences immunological functions. Although ART reduced pulmonary TB pathology, it did not prevent extrapulmonary spread of TB.

## Results

### SIV/ART/TB animals developed similar signs of TB disease as SIV/TB with immune activation

To address the impact of ART controlled viral suppression on SIV/Mtb co-infection, the NHP were randomized into 4 groups: TB-only, SIV/TB, SIV/ART/TB, and SIV-only. SIV infection occurred for 16 weeks prior to Mtb infection with a subset of animals initiated on ART treatment at day 3 of SIV infection, based on prior studies in which viral replication was suppressed despite established viral tissue reservoirs (30) (Figure 1A). Subsequent Mtb infection was planned for 12 weeks in all relevant groups. Serial PET CT imaging, clinical, immunologic and bacterial outcomes were compared at necropsy. We focused our comparisons between SIV/TB and SIV/ART/TB groups with secondary analyses including TB- and SIV-only groups.

**Figure 1.**
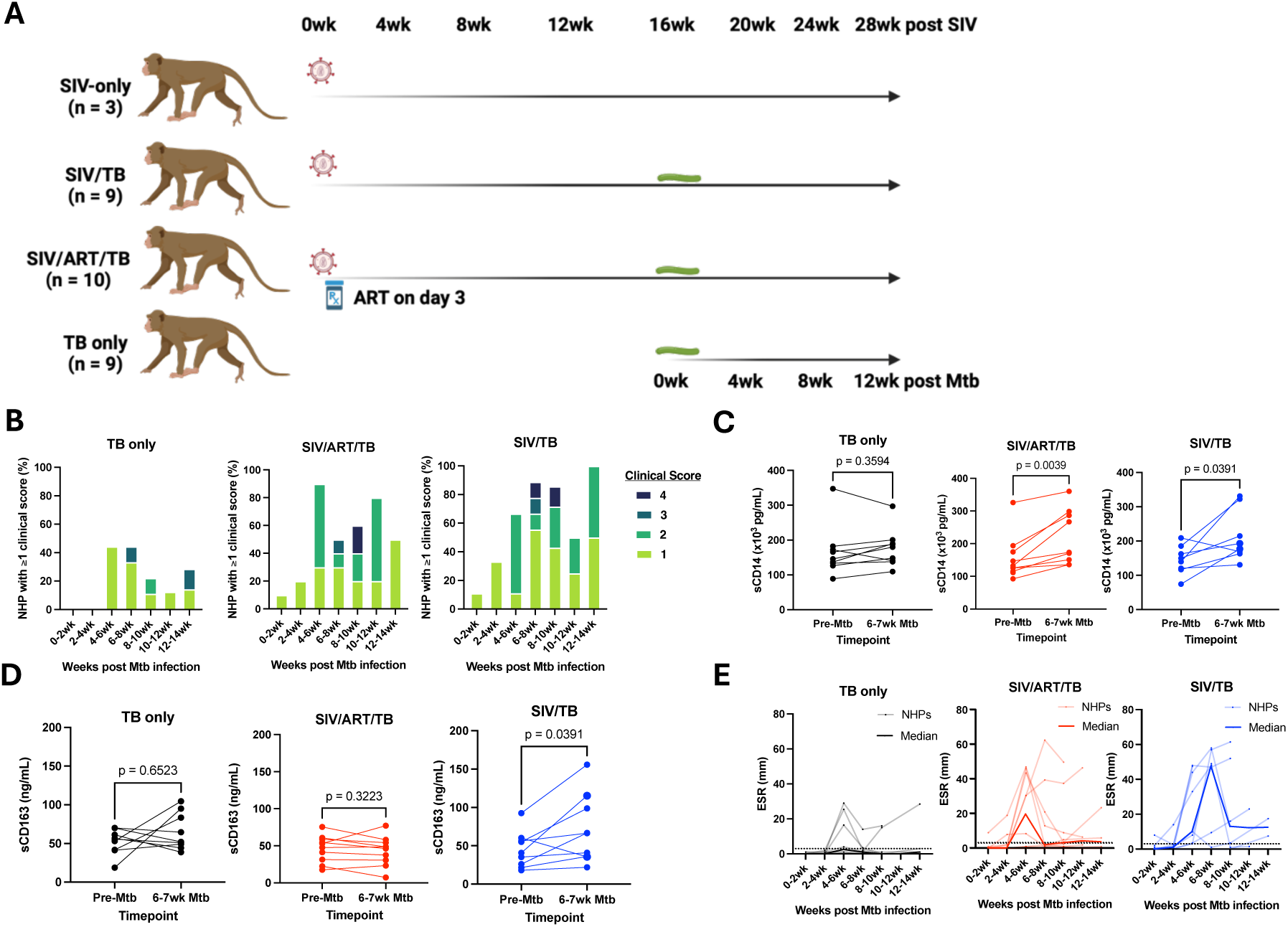
A) Study design and markers of disease. Adult cynomolgus macaques were randomized to receive either SIV infection alone (n=3), SIV infection for 16 weeks and then M. tuberculosis (Mtb) challenge (n=9), SIV with antiretroviral treatment (ART) for 16 weeks and then Mtb challenge (n=10), or Mtb infection alone (n=9). ART was continued throughout the course of SIV and Mtb infection. Mtb infection (low dose Erdman) progressed for 12 weeks with serial blood, airway and lymph node sampling and PET CT performed. B) Clinical scores within each treatment group are shown over the course of Mtb infection. C and D) Interval change in immune activation markers, sCD14 and sCD163, are shown by group. Limited of detection for sCD14 is 125 pcg/ml and sCD163 is 0.469 ng/ml. E) Erythrocyte sedimentation rates (ESR) during the course of infection by experimental group are shown. Each light-colored line represents a single animal over time; dark lines represent median at each timepoint. Normal range is 0-2mm. Wilcoxon matched-pairs signed rank test used for analysis in B and C. TB only (n=9), SIV/ART/TB (n=10), SIV/TB (n=8-9).

Animals randomized to SIV infection without ART showed substantial reductions of CD4 T cells in blood, airways, and tissues (Supplemental Figure 1). Plasma viremia peaked at 1-week post-infection and reached a setpoint by approximately 4 weeks (Supplemental Figure 1A). The level of plasma SIV RNA among SIV/TB NHP was similar to the SIV-only animals. Sustained suppression of SIV replication occurred by 3 weeks of ART in the SIV/ART/TB NHP. This effectively prevented the loss of CD4 T cells within multiple compartments resulting in significantly higher CD4 T cells counts in the blood, airway, and peripheral lymph nodes compared to SIV/TB NHP across all timepoints pre- and post-Mtb infection (Supplemental Figure 1). SIV-only and SIV/TB NHP also maintained similar CD4 T cell frequencies across all time points both before and during Mtb infection in blood, airway, and peripheral lymph nodes. Reductions in CD4 T cells in the SIV only and SIV/TB NHPs were associated with higher CD8 T cell counts (Supplemental Figure 1,2).

**Figure 2.**
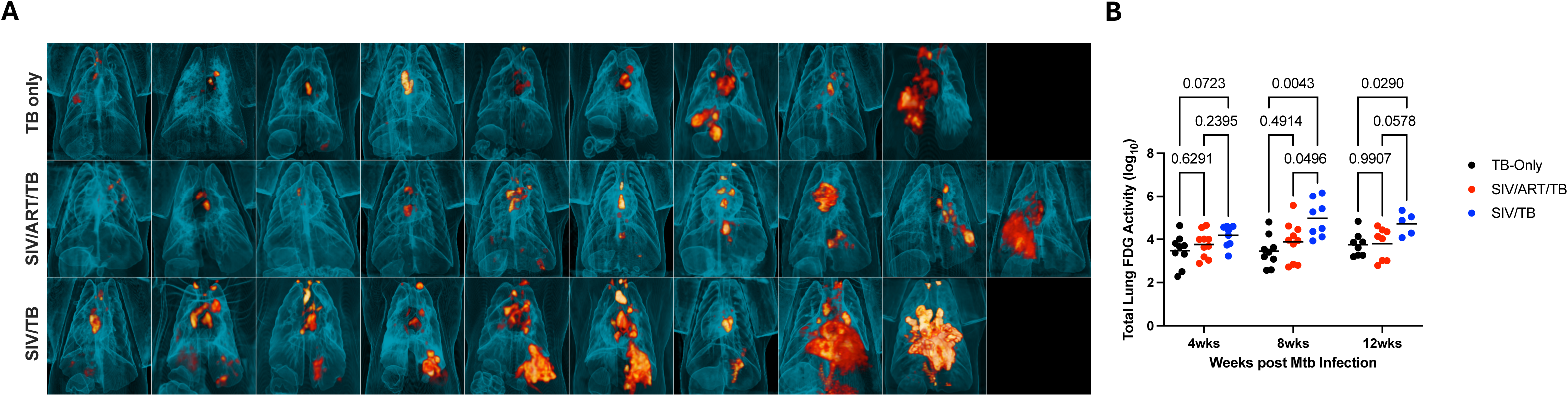
PET CT images of tuberculosis-involved lung and thoracic LN at necropsy. A) Top row: TB Only; Middle row: SIV/ART/TB; Bottom Row: SIV/TB. Images are arranged in each group by total CFU (lowest CFU of the group on the left and highest on the right). B) Total lung FDG activity is presented at 4, 8, and 12wks post Mtb infection for each cohort. A mixed-effects analysis with Tukey’s multiple comparison test were used to determine significance (P > 0.05).

After Mtb challenge, both SIV/TB and SIV/ART/TB macaques displayed more clinical and immunologic signs of early Mtb infection than animals with Mtb infection alone (Figure 1B-E). To quantify and compare clinical signs of disease, we modified our previously published clinical scoring system (31) to numerically quantify signs of TB disease (see methods) (Figure 1B, Supplemental Figure 3). Peaking at 4-6 weeks post-Mtb infection, 40% of the TB-only NHPs developed at least 1 sign of TB similar to previously published studies (26). Yet 60-90% of animals in the SIV/ART/TB and SIV/TB groups had at least one sign of disease at 4-6 weeks that continued throughout infection, often with both microbiologic markers and clinical signs of disease (Figure 1B, Supplemental Figure 3). We measured markers of immune activation, soluble CD14 (sCD14) (macrophage activation marker) and sCD163 (monocyte/macrophage scavenger receptor), often observed in humans with HIV/Mtb co-infection (32). Both the SIV/ART/TB and SIV/TB NHP had significant elevations in sCD14 after Mtb infection but only the SIV/TB group had increased levels of sCD163 (Figure 1C,D). Erythrocyte sedimentation rate (ESR), a marker of systemic inflammation, can be increased transiently during acute Mtb or SIV infection and remain elevated with worsening TB disease (26). In the TB only group, the median ESRs were normal throughout Mtb infection, although there were increases in some animals (27). Conversely, SIV/TB NHP maintained high ESRs starting at 4-6 weeks after Mtb infection (median =10mm IQR^25-75^, 3.25-45.75, n = 9), peaking at 6-8 weeks (47.5mm, 3.125-55, n = 8) and many NHPs had elevated ESR thereafter (Figure 1E). SIV/ART/TB animals experienced high ESR at 4-6 weeks after Mtb infection (19.5mm, 0.875-44.13, n = 10) and remained slightly above normal elevated thereafter. In contrast, plasma cytokines levels of Th1 cytokines (e.g., IL-12, IL-2) were generally higher before and during early Mtb infection among SIV/TB animals (Supplemental Figure 4). None of the TB-only and SIV/ART/TB animals reached humane endpoints prior to their predetermined time of necropsy post Mtb challenge (12 weeks). In contrast, the SIV/TB animals median time to necropsy was 8.9 weeks (Supplemental Figure 5).

### ART did not prevent extrapulmonary dissemination of TB among SIV/ART/TB macaques

At necropsy, inflammation in the lungs and thoracic lymph nodes appeared greater in SIV/TB animals (Figure 2A). PET CT-identified lung granulomas and other sites of disease progression were tracked during the course of Mtb infection. Greater total lung inflammation (FDG activity) is noted in the SIV/TB groups at 8 and 12 weeks after Mtb infection (Figure 2B). At necropsy, SIV/TB NHP had greater overall TB pathology, especially in the lungs and extrapulmonary sites, compared to TB only NHP (Figure 3A). However, SIV/ART/TB NHP had greater extrapulmonary involvement than the TB only group (Figure 3B). A higher proportion of SIV/TB NHP developed TB pneumonia (indicating severe disease) compared to the TB only and SIV/ART/TB NHP groups (78% vs 20% vs 30%, respectively, p = 0.03) (Supplemental Figure 6). Total bacterial burden (including lung and thoracic lymph node) among SIV/TB animals was higher than the TB only and SIV/ART/TB groups (Figure 3C). Individual Mtb bacterial growth per lung granuloma varied across groups (Supplemental Figure 6). SIV/TB animals had higher CFU per lung granuloma (and total live and killed Mtb measured by chromosomal equivalents) indicating increased growth and/or reduced killing than the SIV/ART/TB and TB only groups (Figure 4). Higher Mtb growth was also present in thoracic lymph nodes (with and without granulomas being present) in SIV/TB NHP compared to SIV/ART/TB and TB-only NHP (Figure 4). While it is not surprising that SIV infection increases overall pathology and Mtb growth within NHP, early control of SIV replication with ART appeared to prevent disease progression in the lungs and lymph node but did not ameliorate the dissemination of Mtb to extrapulmonary sites.

**Figure 3.**
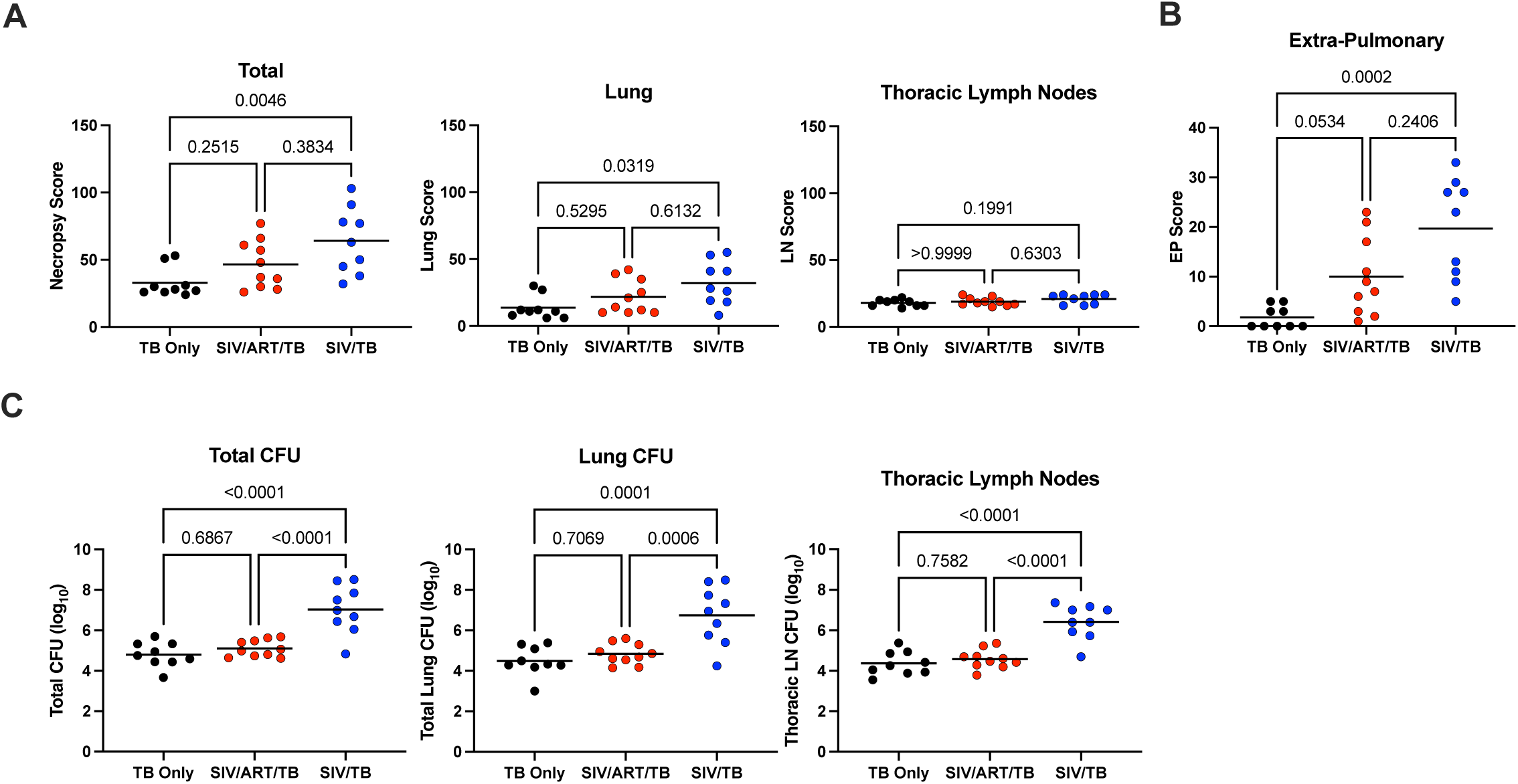
Gross pathology and Mtb burden at necropsy. A) Tuberculosis-associated gross pathology is estimated using a quantitative score at necropsy (necropsy score) that includes disease specific to the lung, lymph node and extrapulmonary sites. B) Tuberculosis-associated extrapulmonary gross pathology and Mtb growth from these sites is used to estimate extrapulmonary score for each animal at necropsy. C) Total Bacterial burden, lung bacterial burden and thoracic LN burden are shown across experimental groups. Lines represent means; each dot represents an animal. Kruskal-Wallis test with Dunn’s multiple comparison adjusted p-values reported in A and B. One-way ANOVA with Tukey’s multiple comparison adjusted p-values reported in C. TB only (n=9), SIV/ART/TB (n=10), SIV/TB (n=9).

**Figure 4.**
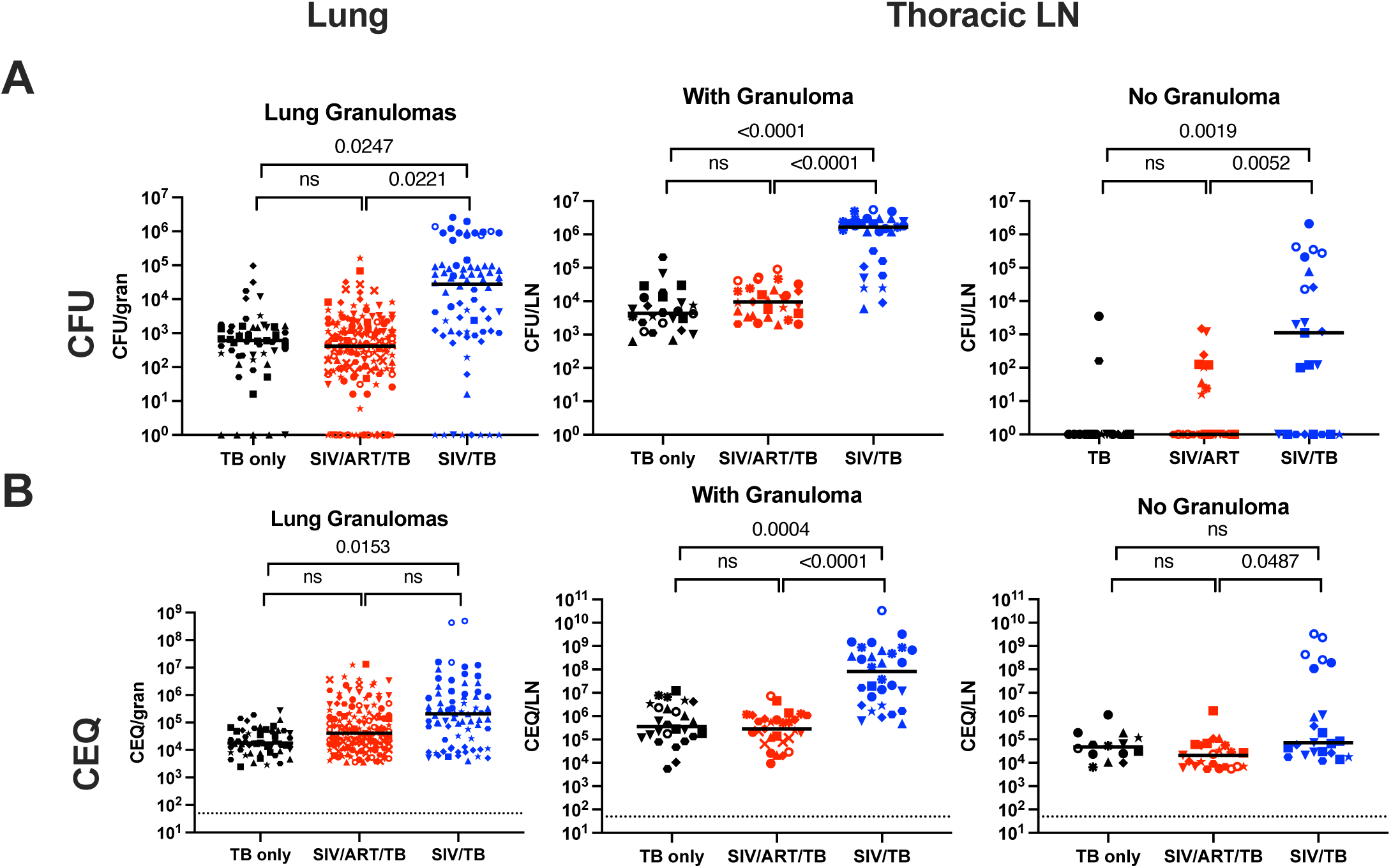
Comparison of viable and total (viable and killed) Mtb per granuloma by group. A) Viable Mtb growth (CFU) by experimental group in lung granulomas, thoracic lymph nodes (with and without granuloma) are shown. B) Total Mtb burden includes both viable and killed Mtb that is measured by chromosomal equivalents (CEQ) in lung granulomas and thoracic lymph nodes. Mixed effect model (animal as a random effect and treatment group as a fixed effect) was used; Tukey HSD adjusted p-values (for p < 0.10) reported. “ns” means not significant (p > 0.10).

### SIV-induced T cell changes are not completely ameliorated by ART

Given that peripheral blood responses do not accurately reflect those in the lung, we focused on the immunological responses in lung granulomas (33). Reduced frequency and total CD4 T cells were noted within lung granulomas of SIV/TB NHP but were preserved in SIV/ART/TB NHP (Figure 5). Similarly, the absolute number of HLA-DR expressing CD4 T cells were lower in the SIV/TB NHP compared to the other cohorts (Figure 5). Given the reduction in CD4 T cells in SIV/TB granulomas, it is not surprising that higher frequencies of CD8 T cells were observed in this group compared to others. The frequencies of CD8 T cells producing any Th1 cytokine (TNF, IFN-γ, or IL-2) were higher in the SIV/TB and SIV/ART/TB granulomas compared to TB only NHP, possibly due to increased bacterial burden and stimulation by SIV or Mtb antigens. Greater CD4 T cell expression of CD38 (activation marker and T cell regulator) was observed among the SIV/TB NHP compared to TB only controls (Supplemental Figure 7), which has been observed in blood of HIV/Mtb co-infected humans (34). We also utilized t-SNE to visualize more complex cell populations (Supplemental Figure 8). The most notable visualized patterns include shifts in phenotypic character among the CD4 and CD8 populations between the TB only and the SIV (+/− ART) groups. These data suggest that while early ART restores some SIV-induced immunological changes, T cell functions are still altered compared to the TB only NHP.

**Figure 5.**
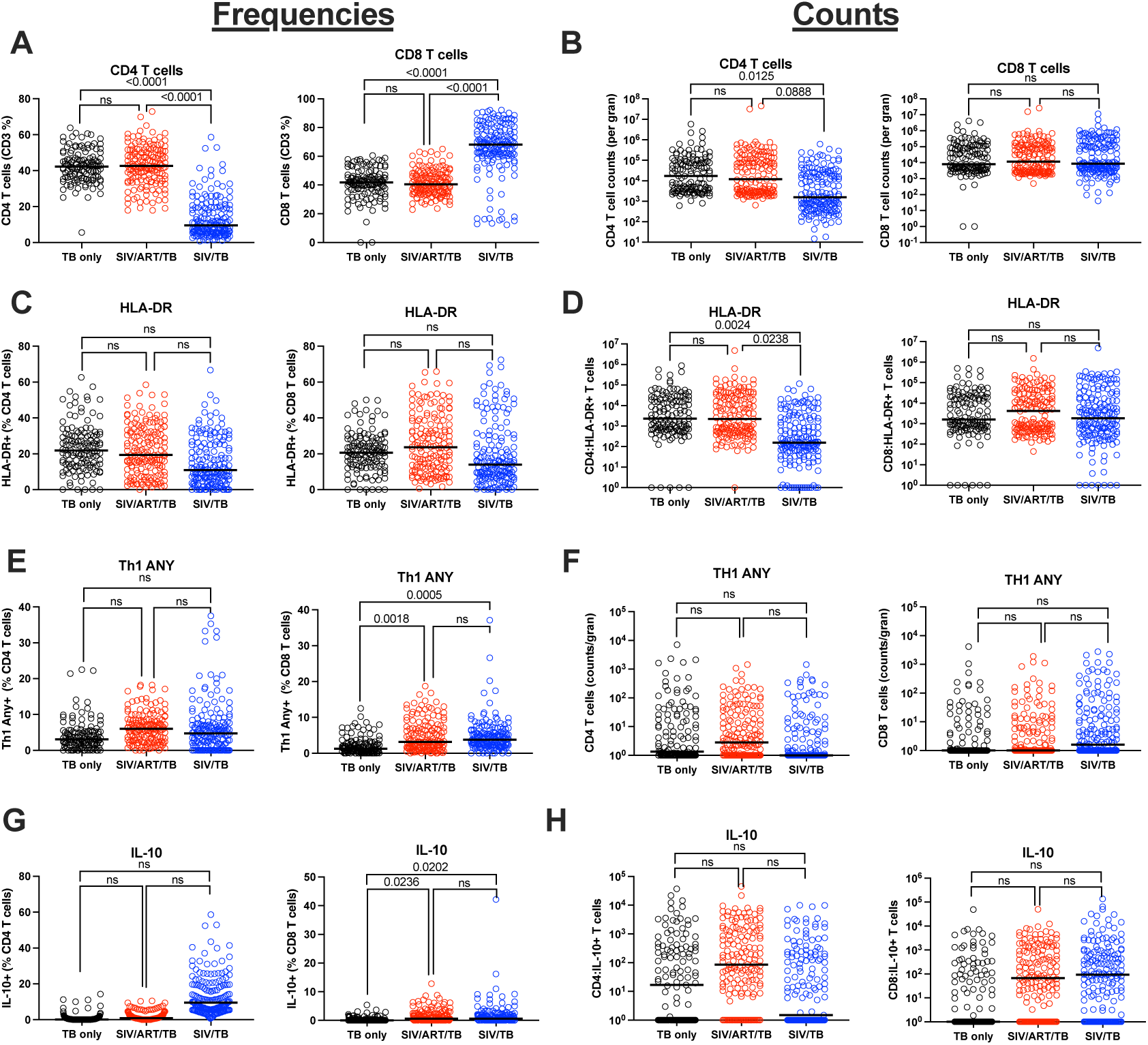
Immunophenotyping of granulomas by experimental group. A and B) Frequency and absolute numbers of CD4 and CD8 T cells per granuloma are shown. C and D) Frequency and absolute number of HLA-DR expressing CD4 and CD8 T cells within granulomas are shown. E and F) Frequency and absolute numbers of the CD4 and CD8 T cells expressing any Th1 cytokine (at least one of the following: IFN-γ, TNF, or IL-2) within granulomas is shown. G and H) Frequency and absolute numbers of IL-10 expressing CD4 and CD8 T cells is shown. Each circle represents a granuloma; lines are medians. Mixed effect model (animal as a random effect and treatment group as a fixed effect) was used; Tukey HSD (Honestly Significant Difference) adjusted p-values (for p < 0.10) reported. “ns” means not significant (p > 0.10).

Thoracic LYMPH NODEs play an important role in T cell priming that is critical to the adaptive immune response and can harbor both Mtb and SIV (35, 36). While the frequency of CD4 T cells among SIV/TB LYMPH NODES were significantly lower than other groups (and the frequency of CD8 T cells consequently higher), the SIV/ART/TB animals had higher CD4 T cells than the other groups (Supplemental Figure 9). CD4 and CD8 T cells expressing CD38 in thoracic lymph nodes were significantly higher in SIV/TB NHP compared to TB only and SIV/ART/TB NHP. Lastly, greater CD4 T cell expression of the degranulating protein CD107a was observed in the SIV/TB group compared the other groups. Based on t-SNE visualization, changes in surface marker and functional phenotypes were noted in both CD4 and CD8 populations in the lymph nodes (Supplemental Figure 10). Perhaps more dramatically than the lungs, these data indicate that SIV changes immunological responses during TB infection and ART does not completely ameliorate those changes.

### ART did not prevent Mtb-barcode dissemination to extrapulmonary sites

While it was clear that there was greater dissemination in the SIV/TB animals, we were surprised to see increased extrapulmonary dissemination in the SIV/ART/TB animals (Figure 6A), prompting further analysis. We tracked the Mtb barcodes with PET CT images across time points to compare bacterial establishment and dissemination within and across experimental groups (37). There was no difference in the number of unique barcodes established between each cohort (Figure 6B). There was no difference in the number of unique barcodes observed in early lung granulomas (granulomas established at 4 weeks post Mtb challenge by PET CT) among the different cohorts (Figure 6C). Nor were there differences in unique barcode counts in thoracic LYMPH NODES across all groups (Figure 6 D). However, both SIV/ART/TB and SIV/TB animals had greater numbers of barcodes shared between extrapulmonary and thoracic LYMPH NODE sites compared to the TB only NHP (Figure 6E). Given that there are multiple thoracic LYMPH NODES (e.g., bilateral hilar and carinal LYMPH NODES) and extrapulmonary sites (e.g., liver, spleen, kidney), we examined the proportion of tissues in each compartment that shared barcodes. Again, both the SIV/ART/TB and SIV/TB NHPs had greater proportions of tissues sharing barcodes than the TB only animals (Figure 6 F,G). No differences were observed in the overall diversity of barcodes within each animal based on treatment group (Supplemental Figure 11). In short, unexpectedly SIV did not influence the establishment of Mtb infection; we expected a greater number of unique barcodes in SIV/TB NHPs compared to other groups. Thus, while initial Mtb infection in the lungs and thoracic LYMPH NODES was unchanged, SIV infection was associated with greater dissemination to thoracic LYMPH NODEs and extrapulmonary compartments, resulting in greater spread of Mtb independent of ART suppression.

**Figure 6.**
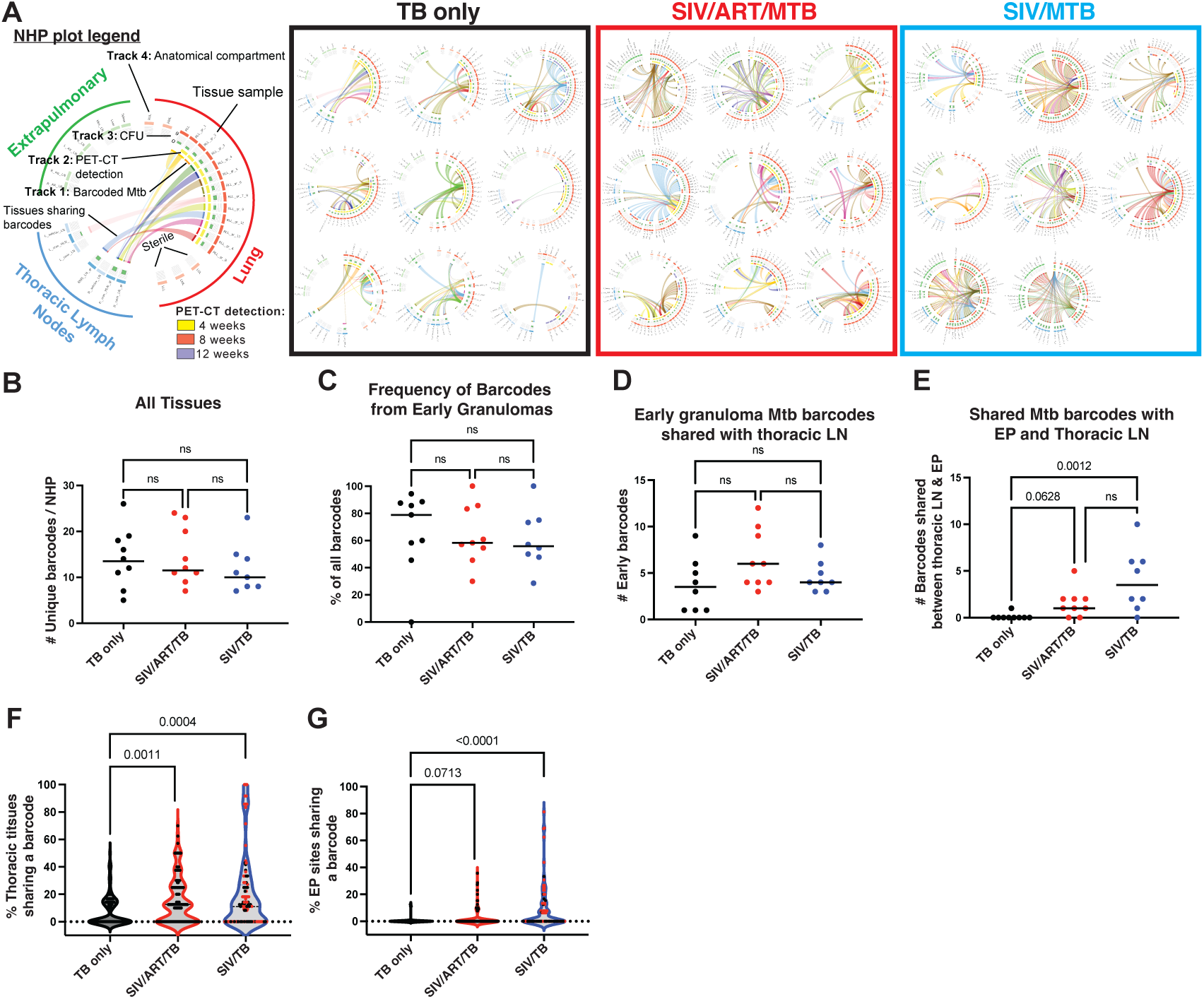
Dissemination and unique Mtb barcode tracking. A) Barcoded Mtb infection of and dissemination across tissues. Circos plots were generated to visualize the barcoded *M. tuberculosis* strains identified in each animal and separated by treatment groups. As described in a legend (top left), each color represents a different barcode sequence identified and barcodes shared across tissues are linked by a ribbon (Track 1). Lung granulomas that were detected by PETCT at 4- (yellow), 8- (red) or 12-weeks (purple) after infection are indicated in Track 2; and the total bacterial burden per tissue sample at necropsy is provided as histograms in Track 3. Sampled tissues are organized by anatomical compartment, including those from lungs (red), thoracic lymph nodes (blue), and extrapulmonary sites (green). Lung sites are also grouped by lung lobe: RUL, right upper lobe; RML, right middle lobe; RLL, right lower lobe; LUL, left upper lobe; LML, left middle lobe; LLL, left lower lobe; Acc, accessory lobe. Sites in lighter shading represent sampled tissues that were found to be sterile. B) Number unique barcodes in each animal (from all sampled tissues). C) Frequency of barcodes from granulomas seen granulomas at 4 weeks after Mtb infection (early). D) Number of barcodes from thoracic lymph nodes that are shared with barcodes from early granulomas. E) Number of barcodes shared between extrapulmonary sites and thoracic lymph nodes. C – E) Each dot represents an animal; lines represent medians. TB only (n=8-9), SIV/ART/TB (n=9), SIV/TB (n=8). F) Proportion of thoracic LN tissues that share barcodes. G) Proportion of extrapulmonary sites that share barcodes. F – G) Each dot represents a barcode; red dots indicate barcodes from animals that did not make it to the planned end of study. B – G) Kruskal-Wallis test with Dunn’s multiple comparison adjusted p-values reported (“ns”: not significant).

## Discussion

In this study, we aimed to elucidate the influence of aggressive ART treatment during SIV/Mtb co-infection on Mtb disease pathogenesis using the NHP model. ART displayed remarkable efficacy in controlling SIV replication and preventing severe pathology associated with SIV/Mtb co-infection. However, it remained ineffective in mitigating bacterial dissemination to extrapulmonary sites. There was a noticeable convergence in clinical symptoms of TB between SIV/TB and SIV/ART/TB groups that we speculate is due to the modified immunological response, including immune activation as seen by sCD14, in the ART group. Both SIV/TB and SIV/ART/TB NHP cohorts had elevated systemic inflammatory markers (ESR), early detection of Mtb growth from gastric aspirates or BAL, and other signs of systemic illness. These clinical manifestations of TB are correlated with increased TB pathology (21, 38, 39). Not surprisingly, CD4 T cells from SIV/TB NHPs were significantly lower in the blood, airways, and granulomas compared with other groups. The reduced number of HLA-DR+ CD4 T cells observed within the SIV/TB NHP is consistent with the HIV literature in which HIV exhibits a preference for infecting HLA-DR+ CD4 T cells (40). While early ART treatment prevented the loss of CD4 T cells, the functional profiles of both CD4 and CD8 T cells in lung granulomas and thoracic LYMPH NODES were distinct from TB only controls.

These studies provide new insights into Mtb dissemination during SIV/Mtb co-infection. Remarkably, primary SIV infection did not increase the establishment of Mtb infection, as the number of unique barcodes was similar across groups. Similar proportions of barcodes were found in the early granulomas and thoracic lymph nodes (the first sites of Mtb dissemination). However, SIV/TB and SIV/ART/TB NHPs had more unique barcodes shared between extrapulmonary and thoracic LYMPH NODE sites and greater proportions of tissues sharing these barcodes within the thoracic LYMPH NODES and extrapulmonary sites compared to TB only (Figure 6); these findings support increased dissemination in SIV-infected macaques regardless of early and sustained ART. Notably, others have reported that the lymphatic system likely plays a pivotal role in Mtb dissemination (41) and extrapulmonary TB is associated with disruptions in LYMPH NODE pathology (42). This increased extrapulmonary dissemination without overt increased Mtb growth in thoracic LYMPH NODES and lung suggests that Mtb dissemination may be driven by factors beyond bacterial load alone. Notably, the comparable Mtb growth in lung and thoracic LYMPH NODE compartments points to a previously unrecognized mechanism of SIV-mediated dissemination that persists despite ART intervention. We hypothesize that once Mtb seeds the thoracic LYMPH NODE or extrapulmonary sites, SIV facilitates greater spread of Mtb within those specific anatomic compartments which can result in increased bacterial burden. These findings raise important conceptual questions about the immune factors required to prevent dissemination from specific compartments (e.g., lung, lymph node), particularly as they relate to extrapulmonary spread of Mtb.

Little is known about the pathogenesis of extrapulmonary disease, though it is often correlated with an increase in mortality especially in co-infected individuals (43). PLWHIV are more likely to develop extrapulmonary TB than HIV uninfected individuals (3, 43–45), especially those with low peripheral CD4 T cells (6, 46). The widespread adoption of ART has resulted in a significant reductions in miliary and disseminated TB (47), although other forms of extrapulmonary TB (e.g., lymphadenitis, meningitis) remain(48). In fact, PLWHIV undergoing ART who subsequently develop TB appear to be at a heightened risk of extrapulmonary TB (49). In those individuals already on ART at the time of TB diagnosis, the median duration until the diagnosis was approximately 15 weeks (49), a timeframe mirroring the observations in our current study where all NHP were necropsied between weeks 8 and 13. We would hypothesize that the immune changes that occur with ART, like HIV-immune reconstitution inflammatory syndrome (19), may alter clinical signs and symptoms prompting the evaluation of extrapulmonary TB similar to the “unmasking TB” scenario where a more robust immune response results in more clinical manifestations of existing disease that was previously asymptomatic.

While these studies reveal significant insights into the complexities of HIV/Mtb co-infection, there are limitations to our studies. Given limited resources, the numbers of animals in each group were powered based on total bacterial burden, so our secondary analyses which included more subtle signs to assess factors that might protect against extrapulmonary disease were insufficiently powered. In our case, the duration of SIV infection was 4 months rather than the years that likely occur in humans. Furthermore, ART began at day 3 after SIV infection, which could be considered an unusual clinical scenario in humans, where HIV ART is often initiated months to years after HIV infection. However in clinical practice, there are uncertainties in the precise timing of HIV and Mtb infections, the duration of these co-infections, and adherence to ART. Lastly, the rate of the extrapulmonary TB disease in humans is likely severely underestimated given the lack of diagnostic tools to adequately assess it (42)

Certainly ART has long term protective effects among PLWHIV (3) and early initiation of ART can reduce the risk of TB (50). The surprising rates of extrapulmonary disease after ART treatment and the alterations in T cell populations noted in the lung and lymph nodes underscore the need to understand the immunologic mechanisms of TB progression and how ART modulates immune responses impaired by HIV. Further studies are needed to develop better diagnostic and therapeutic strategies in HIV/Mtb co-infected individuals.

## Methods

### Sex as a biological variable

Our study examined both 28 male and 4 female animals (Supplemental Table 1). More male nonhuman primates were utilized in this study due to availability.

### Animals

Adult cynomolgus macaques (*Macaca fascicularis*) (>4 years) (Valley Biosystems, Sacramento, CA) were screened for other co-morbidities (e.g., parasites, SIV, Mtb) and then randomly assigned into 4 different experimental groups: TB-only (n=10), SIV/ART/TB (n=10), SIV/TB (N=9), and SIV only (N=3) (Supplemental Table 1). A sample size of 10 animals per treatment group was calculated to detect a difference in lung inflammation of log_10_1.1 with 80% power (alpha=0.05). The majority of animals were male due to the limited availability of females. To limit any gender bias, we attempted to ensure that each experimental group had an equal proportion of females across groups. SIV-designated macaques were infected with 7×10^4^ IU of SIV_B670_ (ARP-633, HIV Reagent Program) via intravenous injection. SIV_B670_ infection was confirmed by detection of plasma SIV RNA. Three days post SIV infection, SIV/ART/TB designated macaques were treated with tenofovir (20 mg/kg/dose, TFV) and emtricitabine (40 mg/kg/day, FTC) plus dolutegravir (2.5 mg/kg/day, DTG) given via subcutaneous route daily (30). After 4 months of SIV infection, macaques were infected with low dose (8-17 CFU per monkey) of digitally barcoded *M. tuberculosis* (Erdman strain (37)) via bronchoscopic instillation to the lower lung lobe and housed in a Biosafety Level 3 (BSL-3) NHP facility, as previously described (21, 22, 27) (Figure 1A). Mtb infection was confirmed by the detection of TB-specific lesions on serial Positron Emission Tomography using 18-flourodeoxyglucose (FDG) probe and Computed Tomography (PET CT) scans and confirmed at necropsy. Blood was obtained every 1-4 weeks throughout the course of the study for serial SIV viral load and systemic immune responses, as previously described (21). Bronchoalveolar lavage (BAL) was performed monthly throughout the study. Peripheral (axillary and inguinal) lymph node biopsies were performed prior to SIV infection, 4 weeks post SIV infection, prior to Mtb infection, and 4 weeks after Mtb infection.

### Study approval

All animal procedures were approved by the University of Pittsburgh Institutional Animal Care and Use Committee in compliance with the Animal Welfare Act and Guide for the Care and Use of the Laboratory Animals.

### Clinical measurements and scoring

Serial clinical, microbiologic, radiographic and immunologic metrics were regularly obtained. After Mtb infection, BAL and gastric aspirate (GA) were performed every 4 weeks for Mtb growth and ESR every 2-4 weeks. Daily clinical assessments were conducted and animals that showed evidence of clinical deterioration were monitored and euthanized prior to 12 weeks if they met clinical criteria (i.e., loss of appetite, weight loss, increase in respiratory rate or effort, hunched posture, and dehydration).

Clinical scores were generated by measuring TB specific signs observed from each animal after Mtb infection in 2-week time intervals (0-2 weeks, 2-4 weeks, 4-6 weeks, 6-8 weeks, 8-10 weeks, 10-12 weeks, 12-14 weeks) (31). These signs included: Mtb growth from either GA or BAL, abnormal ESR (>2 mm), coughing, loss of appetite, weight loss, increased respiration, hunched posture, and dehydration. NHP who developed 1, 2, 3, or 4 of these clinical signs of disease during a 2-week time frame were identified.

### PET CT Imaging and analysis

To follow in vivo disease progression, serial PET CT imaging was performed using 2-deoxy-2-18F-D-deoxyglucose (FDG) as the probe to identify TB lesions (51). Animal studies were performed in cohorts over several years with 2 different PET CT scanners: PET co-registered with CT in the first cohorts were imaged with a microPET Focus 220 preclinical PET (Siemens Medical Solutions) and a CereTom clinical helical CT (NeuroLogica Corp) (21, 27, 52) while the latter cohorts were imaged using a Mediso MultiScan LFER 150 integrated preclinical PET CT (Mediso, Budapest, Hungary).

Each PET scanner’s sensitivity was calibrated by the respective manufacturer’s recommended procedure, involving scan measurements of known quantities of radioactivity within known volumes of solution. All quantities of tracer, for both the purpose of calibration and scientific measurement, were measured with the same dose calibrator. PET CT scans were performed every four weeks after Mtb infection. As previously described (53), Mtb involvement within the lungs and thoracic lymph nodes was measured using several different parameters such as: count of individual granulomas, total lung FDG activity, number of mediastinal lymph nodes with increased FDG avidity with or without the presence of necrosis, presence of extrapulmonary involvement (e.g., liver lesions), as previously described (21, 54).

### Plasma sCD14 and sCD163 measurements

Levels of sCD14 (R&D Systems, Human sCD14 ELISA, Bio-techne, Minneapolis, MN) and sCD163 (MyBiosource, Monkey sCD163 ELISA, San Diego, CA) were measured by commercial kits. Plasma samples from animals were obtained at serial time points and filtered (0.22 um) prior to freezing (−80°C). Samples were then thawed, diluted (1:100 for sCD163, 1:400 for sCD14) in diluting buffer and run according to manufacturer’s instruction. Samples were run in duplicate. Optimal densities were measured using Spectramax Microplate Reader 190 (Molecular Devices, San Jose, CA) using analysis software (SoftMaxPro 7.1.2). Analysis of the sCD14 and sCD163 standards were transformed using 4 parameter logistic regression and linear regression methods, respectively.

### Necropsy

TB specific gross pathology at the time of necropsy was quantified using a gross pathology score system that accounts for the number and size of granulomas, distribution of disease (e.g., lung lobes, lymph nodes, extrapulmonary sites) previously described (55). PET CT matched lesions were harvested and halved for histopathology and homogenized into single cell suspensions for bacterial burden and killing and immunology as previously described (21, 53).

### *Ex vivo* immunologic assays

Flow cytometry was used to determine phenotypic and functional changes in CD4 and CD8 T cells from PBMC, BAL, and peripheral lymph nodes over time. Cells were stimulated with ESAT6 and CFP10 peptides or SIV Gag-Pol peptide pools Mtb (10ug/ml) in media (RPMI+10%hAB), media alone, or PBDU/ionomycin (BEI Resources, Manassas,VA). PBMC were stimulated for 6hrs, while BAL samples were stimulated for 4hrs in the presence of Brefeldin A. Tissue samples were incubated with media alone and Brefeldin A for 4hrs. Flow cytometry was performed on all stimulated cells after staining with a combination of antibodies (Supplemental Table 2). Data acquisition was performed using an LSR II (BD) and analyzed using FlowJo Software v.10.8 (Treestar Inc, Ashland, OR) using a standardize gating strategy (Supplemental Figure 12).

### SIV RNA isolation and quantification from plasma and tissues

Viral RNA was isolated from plasma as previously described (56). Briefly, plasma was separated from EDTA-treated whole blood and stored at −80° C until use. Plasma was centrifuged at 16,000 × g for 1 hr at 4°C. The viral pellet was resuspended in 5 mM Tris-HCl (pH 8.0) containing 200 μg of proteinase K for 30 min at 55°C, followed by 5.8 M guanidinium isothiocyanate containing 200 μg of glycogen. RNA was precipitated with isopropanol, washed with 70% ethanol and resuspended in 5 mM Tris-HCl (pH 8.0) containing 1 μM dithiothreitol and 1,000 U of an RNase inhibitor (RNasin; ThermoFisher) and stored at −80° C until use.

Tissue specific RNA was isolated from single cell homogenates stored in Trizol LS. After thawing, the homogenate was mixed with 1-bromo-3-chloropropane (MRC Labs) at 10:1 and centrifuged at 14,000 x g for 15 min at 4° C. The upper aqueous phase was removed to a new tube and 240 mg glycogen (Roche) was added. Isopropanol was added and mixed before centrifugation at 21,000 x g for 10 min at room temperature. The pellet was washed with 70% ethanol and centrifuged at 21,000 x g for 5 min at room temperature. After the final wash, the pellet was dried, resuspended in RNase-free water, and stored at −80° C until use.

qRT-PCR was performed as previously reported (57). Briefly, cDNA synthesis was performed on the isolated RNA using SuperScript III First-Strand Synthesis SuperMix (ThermoFisher) following manufacturer’s instructions and using supplied random hexamers. qPCR was performed in duplicate wells using SsoAdvanced Universal Probes Supermix (BioRad). Primers and probes were designed for macaque CD4 (tissues only), as previously described (57) and SIV_B670_ *gag*: Forward 5’- GTCTGCGTCATTTGGTGCATTC-3’, Reverse 5’-CACTAGATGTCTCTGCACTATTTGTTTTG-3’, and Probe FAM-CGCAGAAGAGAAAGTGAAACATACTGAGGAAG-TAMRA. Viremia is reported as SIV RNA copies per ml (plasma) or copies per 10^6^ CD4 RNA copies (tissues).

### Quantification of Mtb viable and total (genomic) bacterial burden

Colony forming units (CFU) were used to estimate bacterial growth within single cell homogenates from each individual site, as previously described (21, 25, 55). The sum of CFU within lymph nodes was considered lymph node burden (“LN CFU”) and was defined as the sum of CFU from all thoracic lymph nodes. We quantified extrapulmonary involvement (EP score) through TB disease in EP sites (e.g., liver, peripancreatic lymph node, paracostal abscess, kidney) for which bacterial growth, gross or microscopic evidence of Mtb involvement are assessed (21, 55). Total bacterial burden includes the sum of CFU from the lymph nodes (mediastinal and extrapulmonary) and lung lesions (e.g., grossly normal lung, granulomas, involved lung, or diaphragm granulomas), as previously described (21).

Mtb DNA extractions and qPCR for estimating chromosomal equivalents (estimated live and dead Mtb) were performed as previously described (21, 36). Chromosomal equivalents (CEQ) were assessed relative to a serially diluted standard curve of *M*. *tuberculosis* genomic DNA using quantitative real-time PCR; efficiency for each run was kept between 90% and 110%. Each sample was analyzed in duplicate on a QuantStudio 6 real-time PCR system (ThermoFisher Scientific, Waltham Massachusetts) with a 96-well block using primers targeting SigF and iTaq Universal SYBR Green Supermix (Bio-Rad, Hercules, CA).

### t-Distributed Stochastic Neighbor Embedding (t-SNE) Analysis

Flow cytometry data of gated lymphocytes from SIV/TB (n = 3), SIV/ART/TB (n = 3), and TB-only (N = 2) NHPs were concatenated into single FCS files in FlowJo 10 software based on individual cohort for lung granulomas and thoracic LN. The number of granulomas included within the concatenation file were 39, 43, and 19 in SIV/TB, SIV/ART/TB, and TB-only NHPs, respectively. The number of thoracic LYMPH NODES included in the concatenated file were 12, 14, and 4 in SIV/TB, SIV/ART/TB, and TB-only NHPs, respectively. The concatenated files were down sampled to 250,000 events per cohort and t-SNE-CUDA visualization was performed in Cytobank. Perplexity and the number of iterations were automatically applied. The antibodies utilized for t-SNE analysis were CD3, CD4, CD8, CD38, HLA-DR, TNF, IFN-g, IL-2, CD107a, IL-17, and IL-10.

### Plasma Cytokine and Chemokine Detection

Plasma cytokine and chemokine measurements from animals were performed using the Meso Scale Discovery (MSD) per V-PLEX NHP Cytokine 24-Plex kit (K15058D-2) packaged instruction. The following were examined: IL-12/IL-23p40, IL-2, IP-10, MCP-4, IL-15, MCP-1, IL-6, MIP-1β, IL-7, IL-8, MIP-1α, GM-CSF, TNF-β, IL-8 (HA), Eotaxin-3, IL-16, IL-5, MDC, IFN-γ, and VEGF. All samples were plated as neat. Results are displayed with https://software.broadinstitute.org/morpheus/.

### Barcode diversity

Diversity of mycobacterial DNA barcodes was quantified using the effective number of species, calculated from the Shannon entropy index(58). The effective number was broken down by anatomical compartment and barcode diversity within lung lobes are reported.

### Barcode determination to track Mtb dissemination and CIRCOS plots

To track Mtb dissemination, digitally barcoded Mtb was used as previously described (37). Each library contained approximately 16,000 unique barcodes and 3 independently generated libraries were combined into a master library to increase the number of unique barcodes, ensuring a <2% chance that a barcode would be represented twice if 20 bacteria are randomly selected (37). Scan matched lesions and other tissues were plated for viable Mtb growth after which purified bacterial DNA was extracted, The DNA was then amplified, indexed using AMPure XP beads (Beckman Coulter, catalogue #A63881) and pooled. Each pool was sequences on an Illumina MiSeq System resulting in ∼ 40 millions reads per library. Sequencing data was further analyzed using a custom Perl script (available by request) for quantifying individual barcodes as previously published (37).

Each plot was then manually adjusted such that links between tissues sharing barcodes were drawn, when possible, to originate from lung sites detected early by PET-CT at 4 weeks post-infection with the highest CFU burden as previously published (59). If no tissue in a barcode dissemination network was identified at 4 weeks post-infection, the tissue with the highest CFU burden was chosen as the origin site. However, other lung sites with the same barcode may also be the source of dissemination between tissues.

### Statistical methods

Data were tested for normality with Shapiro-Wilk test. Multiple Mann-Whitney tests with Holm-Šídák adjusted p-values were used to analyze two groups over time. Wilcoxon matched-pairs signed rank test was used to test differences among paired data points. When testing mean differences among 3 groups, one-way ANOVA with Tukey’s multiple comparison adjustment was used for normally distributed data; otherwise, Kruskal-Wallis test with Dunn’s multiple comparison adjustment was used. Lung inflammation over time was compared across groups using two-way ANOVA with Tukey’s multiple comparison adjustment (for comparing all three groups at each time point). For categorical data, Fisher’s exact test was used to compare distributions among treatment groups. Statistical tests above were performed in GraphPad Prism Mac OSX (Version 10.3.1, GraphPad San Diego, CA). For multiple granulomas and thoracic lymph nodes per animal, random effect models (monkey as the random effect and treatment group as the fixed effect) were fit using the restricted maximum likelihood (REML) method (JMP®Pro 17.2.0). Tukey HSD (Honestly Significant Difference) adjusted p-values are reported for pair-wise multiple comparisons. All statistical tests are two-sided, and significance was established at p ≤ 0.05 and trends p < 0.10.

