## Supplemental Figures and Tables for "Antiretroviral treatment does not prevent extrapulmonary tuberculosis during SIV/Mtb co-infection in macaques"

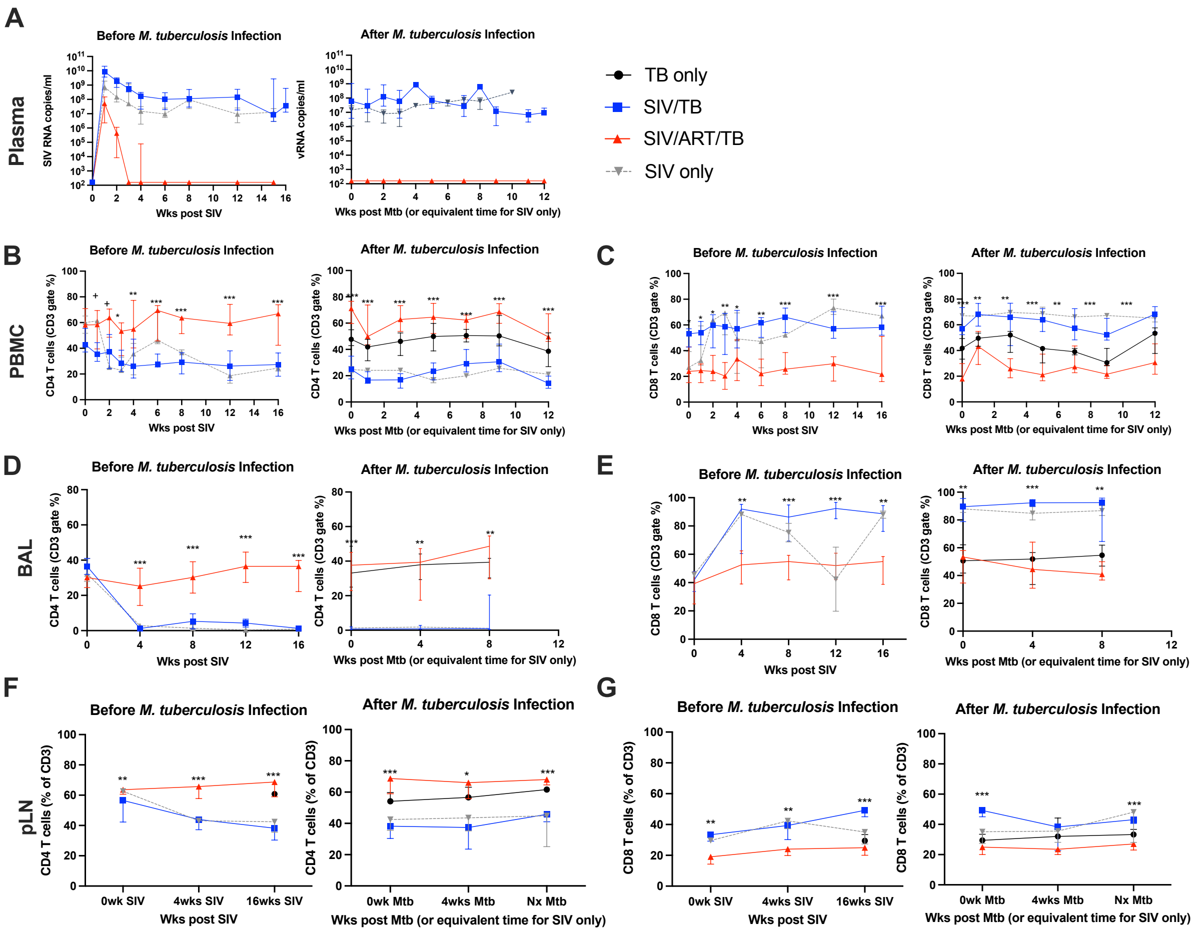


**Supplemental Figure 1.** A) Serial SIV RNA plasma levels before and after M. tuberculosis infection are shown. B and C) Serial frequencies of CD4 and CD8 T cells in the peripheral blood. D and E) Serial CD4 T cells and CD8 T cells in the airway before and after Mtb challenge, F and G) Serial frequencies of CD4 and CD8 T cells in peripheral lymph nodes (pLN). (TB only =9, SIV/TB =9, SIV/ART/TB= 10, SIV only = 3) Median with IQR error bars shown. Statistical analysis was restricted to compare only SIV/ART/TB and SIV/TB groups. Mann-Whitney test run at each time point and adjusted for multiple comparisons by Holm-Šídák method (+: 0.05 < p < 0.10, *: 0.01 < p < 0.05, **: 0.001 < p < 0.01, ***: p < 0.001). Black: TB only; Blue: SIV/TB; Red: SIV/ART/TB; Gray: SIV-only.

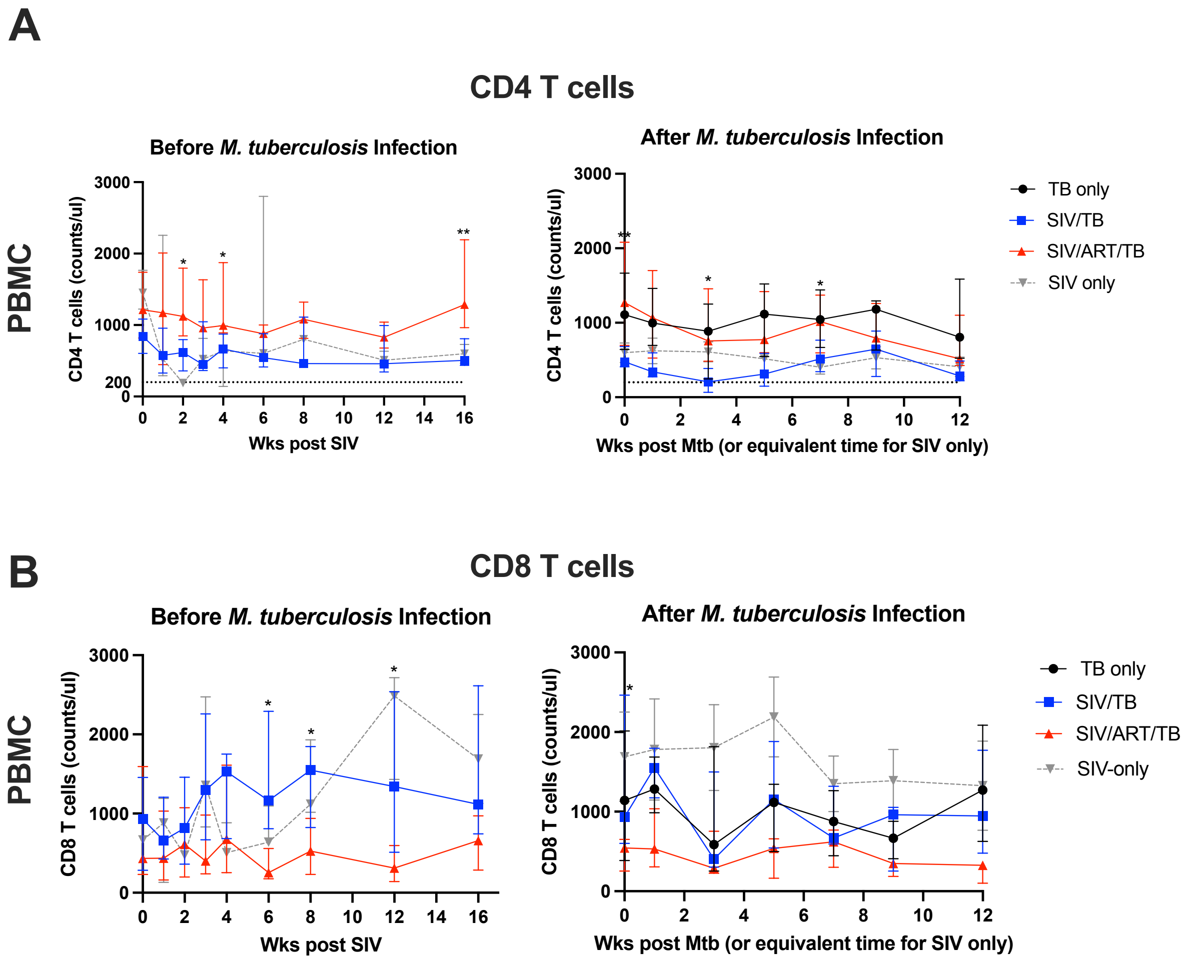


**Supplemental Figure 2.** Absolute numbers of CD4 (A) and CD8 (B) T cell counts in peripheral blood are shown across groups of animals. (TB only = 9, SIV/TB =9, SIV/ART/TB=10, SIV only = 3). Median with IQR error bars shown. Statistical analysis was restricted to compare only SIV/ART/TB and SIV/TB groups. Mann-Whitney test run at each time point and adjusted for multiple comparisons by Holm-Šídák method (+: $0.05 < p < 0.10$, *: $0.01 < p < 0.05$, **: $0.001 < p < 0.01$, ***: $p < 0.001$). Black: TB only; Blue: SIV/TB; Red: SIV/ART/TB; Gray: SIV-only.

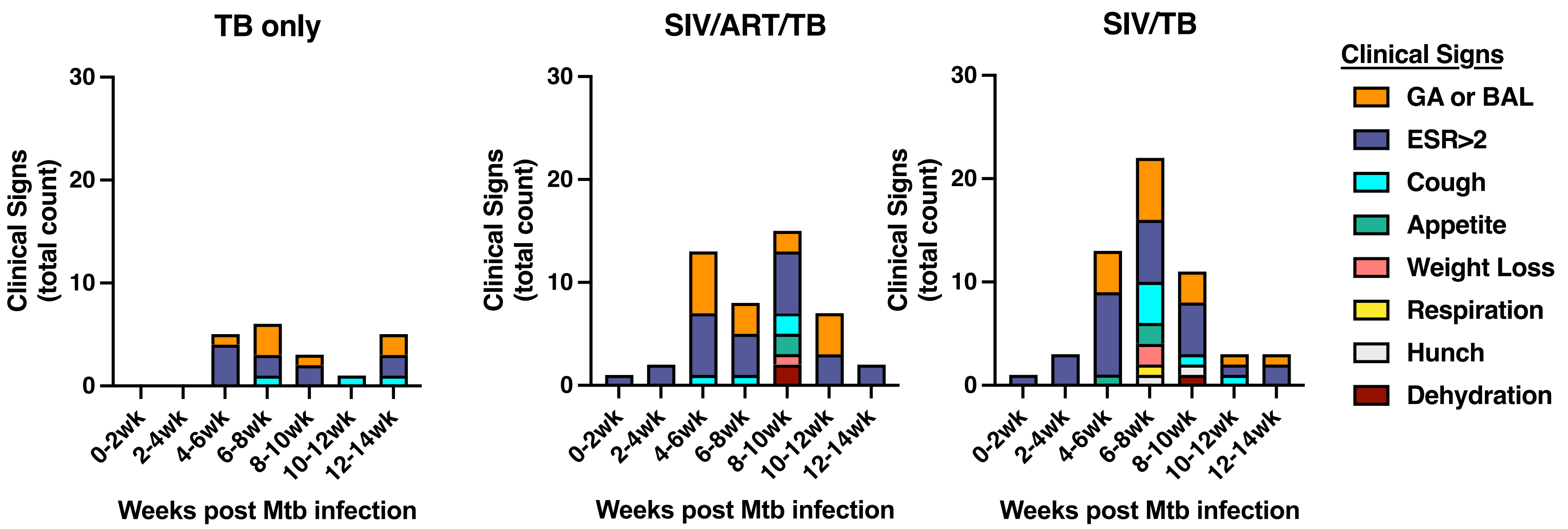


**Supplemental Figure 3.** Detailed clinical signs for each experimental group is shown by stack graph across each time point. Each stack graph represents the sum of clinical signs within each experimental group at that time point. TB only (n=9), SIV/ART/TB (n=10), SIV/TB (n=9).

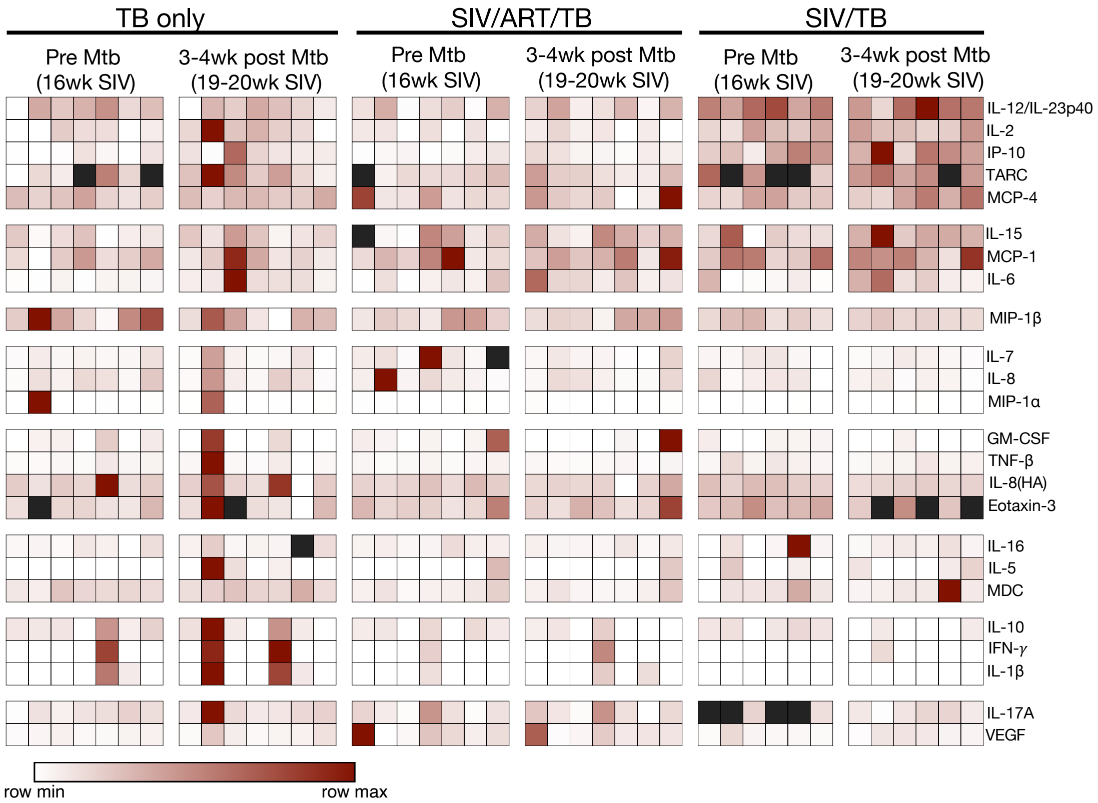


**Supplemental Figure 4. Cytokine and chemokine expression within plasma.** Each column represents a macaque and are divided into their specific infection cohort (TB-only, SIV/ART/TB, and SIV/TB) and time point. Each row represents a different cytokine or chemokine, ranging from smallest (white) to greatest (dark red). The cytokines were grouped using hierarchical clustering on one-minus-Kendall's correlation (and average linkage method). Black squares represent missing data points. This figure was created using Morpheus, https://software.broadinstitute.org/morpheus/.

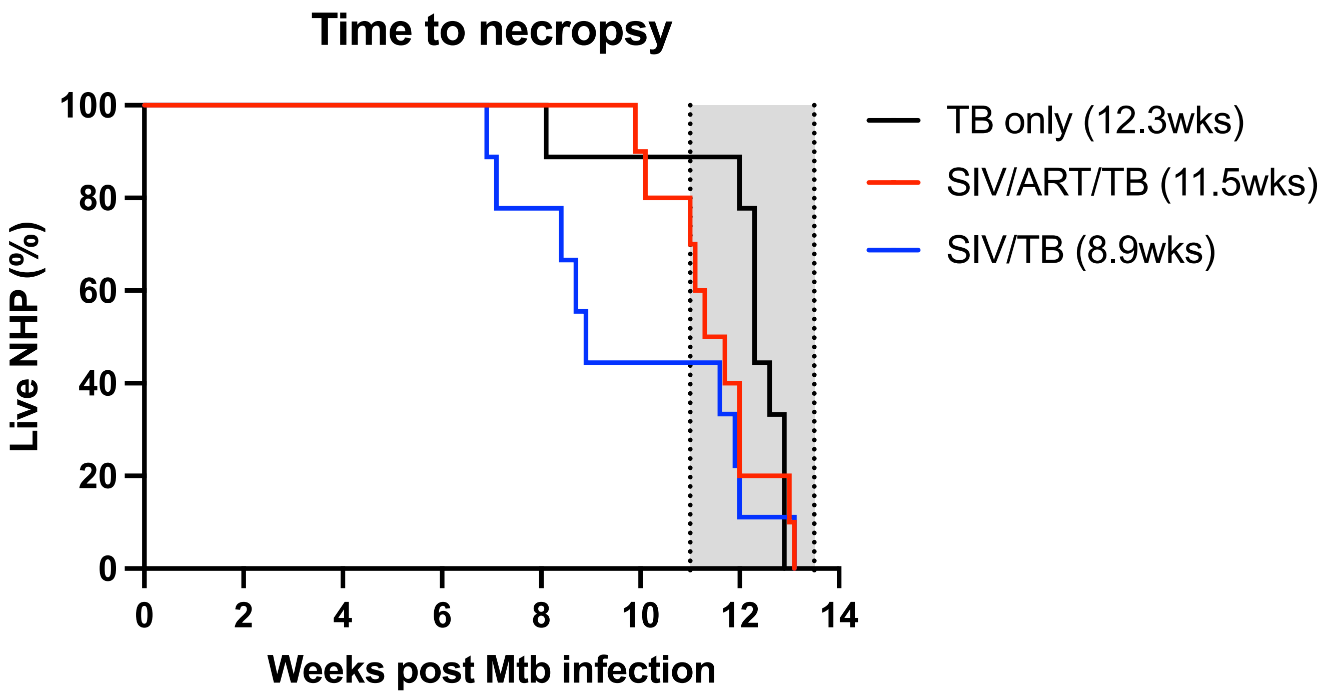


**Supplemental Figure 5**. Proportion of animals surviving to declared endpoint (12 wk) after Mtb challenge. Median weeks of Mtb infection are shown in ( ). Grayed area indicates time of planned necropsy.

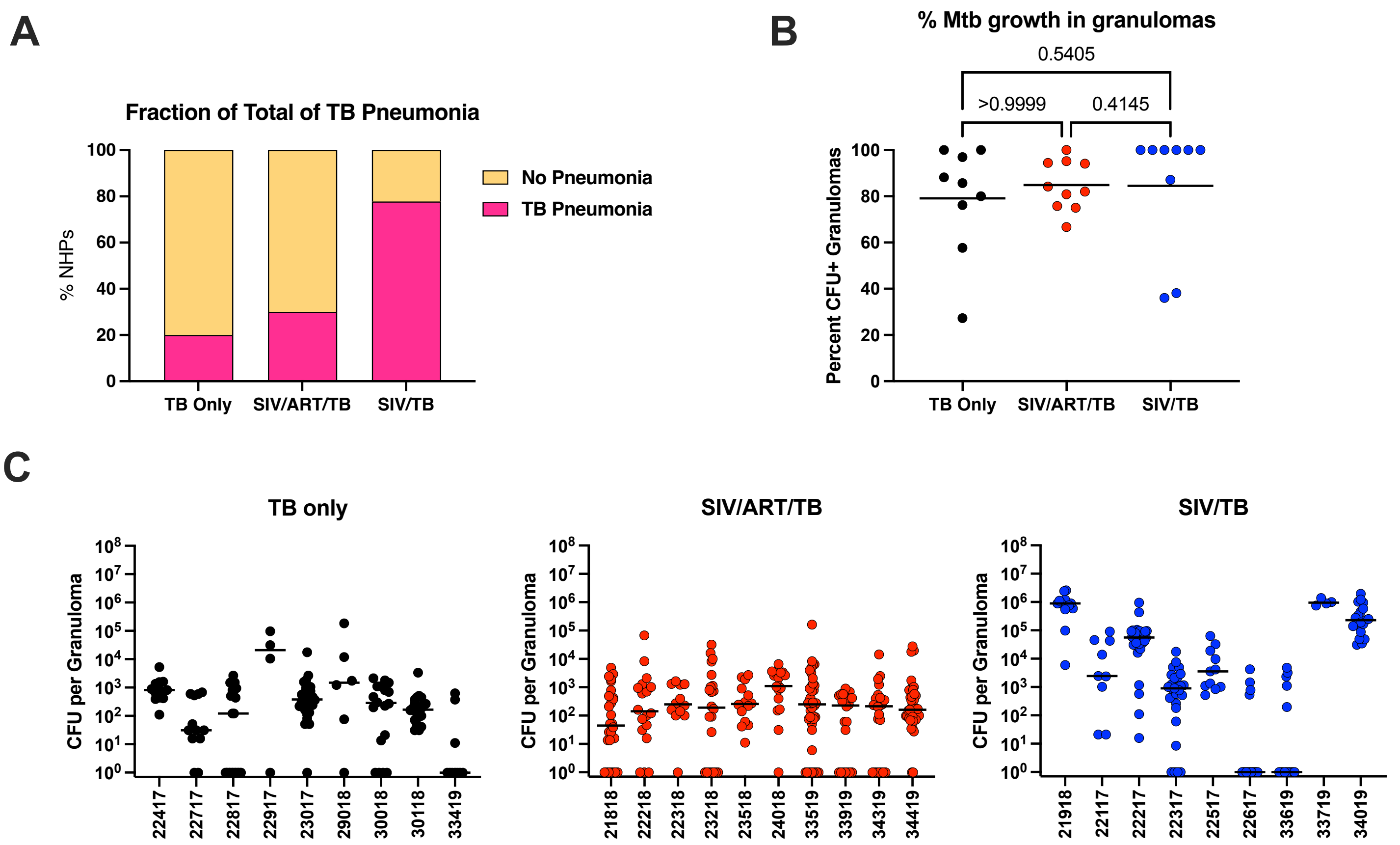


**Supplemental Figure 6**. A) Greater total lung inflammation (FDG activity) is noted among SIV/TB compared to TB only and SIV/ART/TB groups over time at 8 weeks post-Mtb infection. Two-way ANOVA with Tukey's multiple comparisons p-values (<0.05) reported at each time point. B) Proportion of NHPs in each experimental group with TB pneumonia determined by PET CT, Fisher's Exact test p-value 0.0278. C) Percentage of granulomas and clusters with viable Mtb growth. Kruskal-Wallis test with Dunn's multiple comparison adjusted p-values reported. D) Individual bacterial burden (CFU) per granuloma by animal and group.

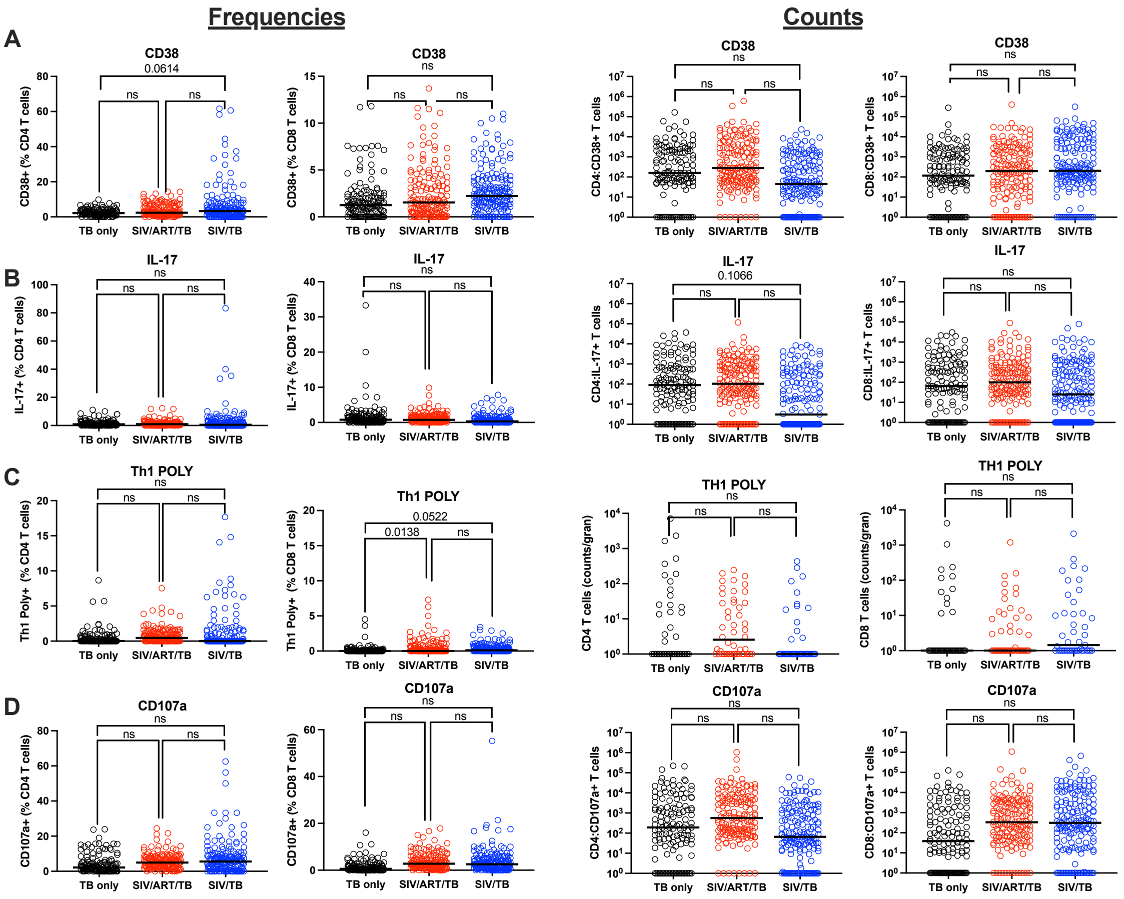


**Supplemental Figure 7**. Immunophenotyping of granuloma specific CD8 and CD4 T cells based on frequencies (left) and absolute cell numbers (right columns). Th1 POLY represents cells that express two or more of the following: IFN-g, TNF, IL-2. Each circle represents a granuloma; lines are medians. Mixed effect model (animal as a random effect and treatment group as a fixed effect) was used; Tukey HSD adjusted p-values (for $p < 0.10$) reported. “ns” means not significant ($p > 0.10$).

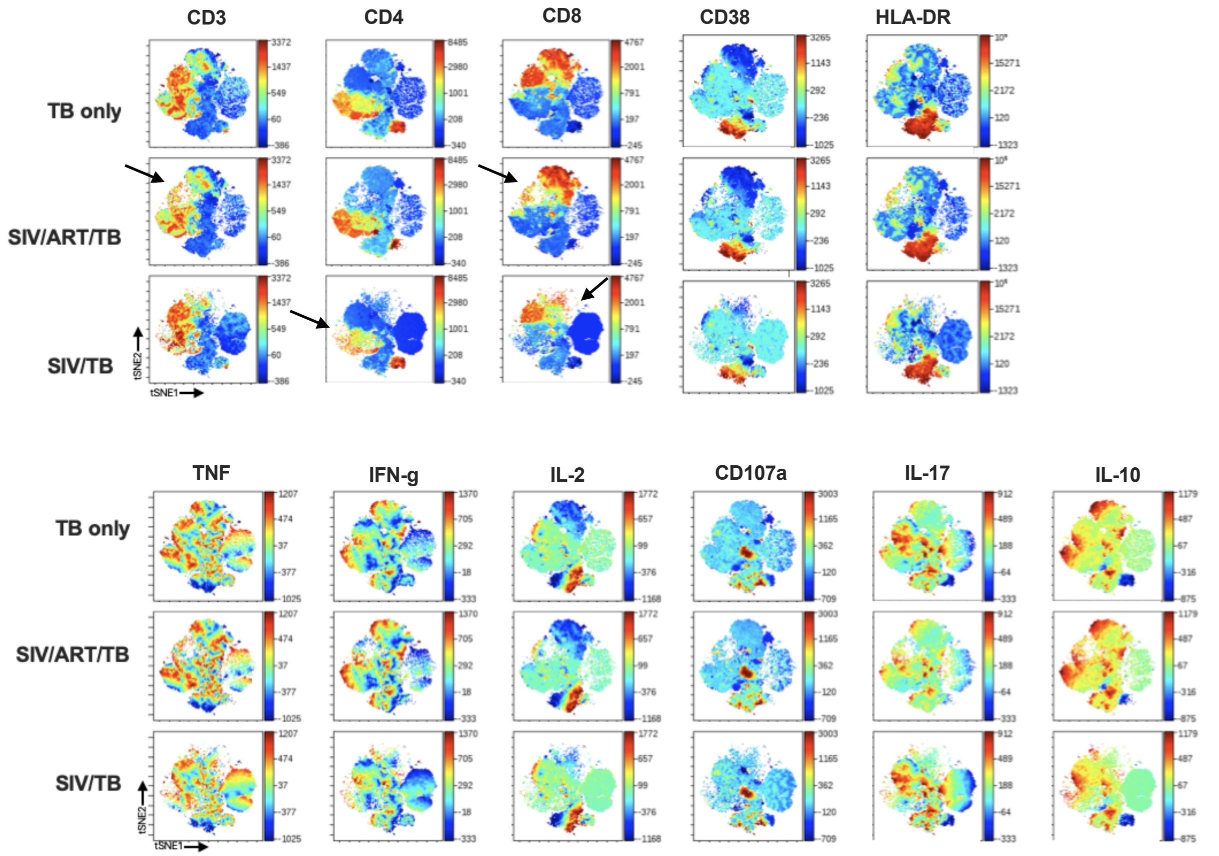


**Supplemental Figure 8**. t-SNE visualization of cellular markers among granuloma-specific lymphocytes. Pseudocolored maps show the relative expression of respective markers where red represents high expression and blue is low expression. Arrows highlight T cell populations based on phenotypic characteristics that differ between experimental groups. (n=39, 43, and 19 lung granulomas from 3 SIV/TB, 3 SIV/ART/TB, and 2 TB-only NHPs, respectively).

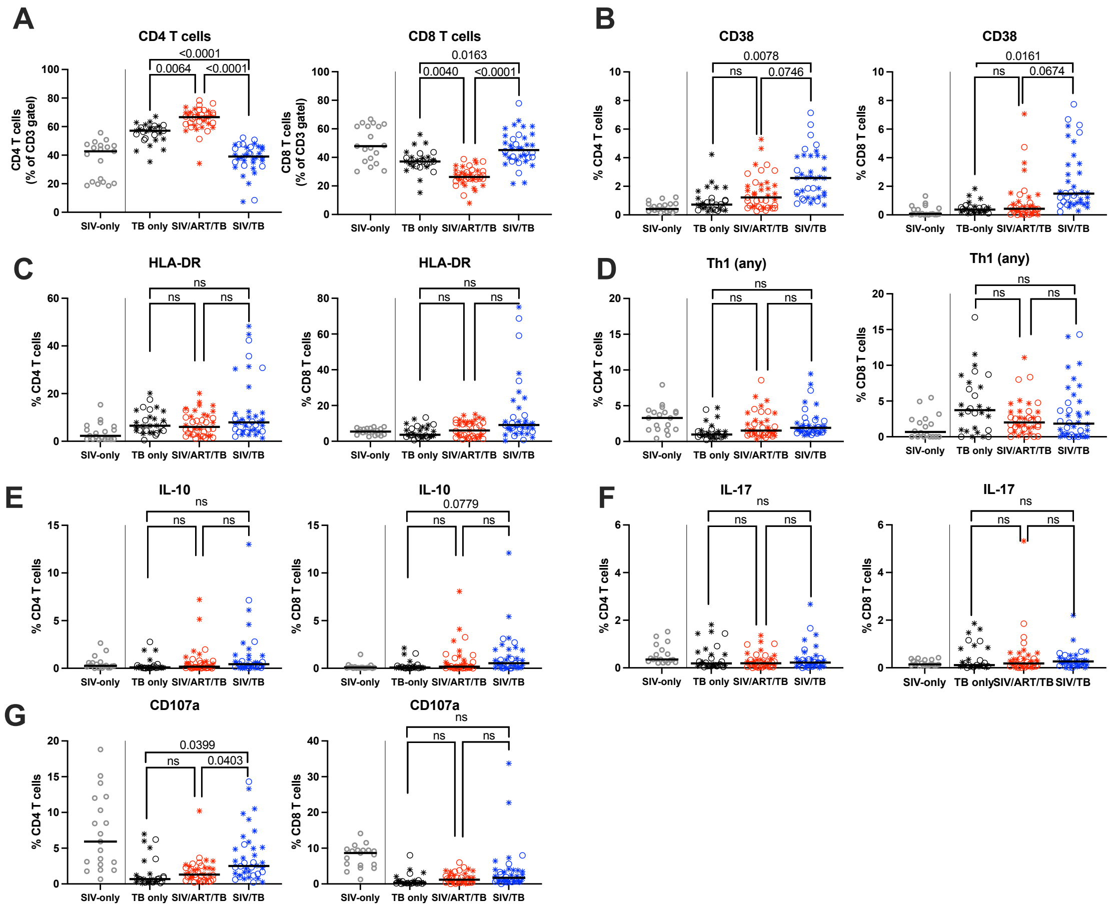


**Supplemental Figure 9.** Frequency and functional characteristics of CD4 and CD8 T cells from thoracic lymph nodes (LN). Th1 (any) represents at least one of the following: IFN-g, TNF, or IL-2. Each circle is a mediastinal LN. Stars represent LN with granuloma. Lines are medians. Mixed effect model (animal as a random effect and treatment group as a fixed effect) was used; Tukey HSD adjusted p-values (for p < 0.10) reported. "ns" means not significant (p > 0.10). SIV only groups is shown for visual representation without statistical comparison. Statistical comparison is performed across TB only, SIV/ART/TB and SIV/TB groups.

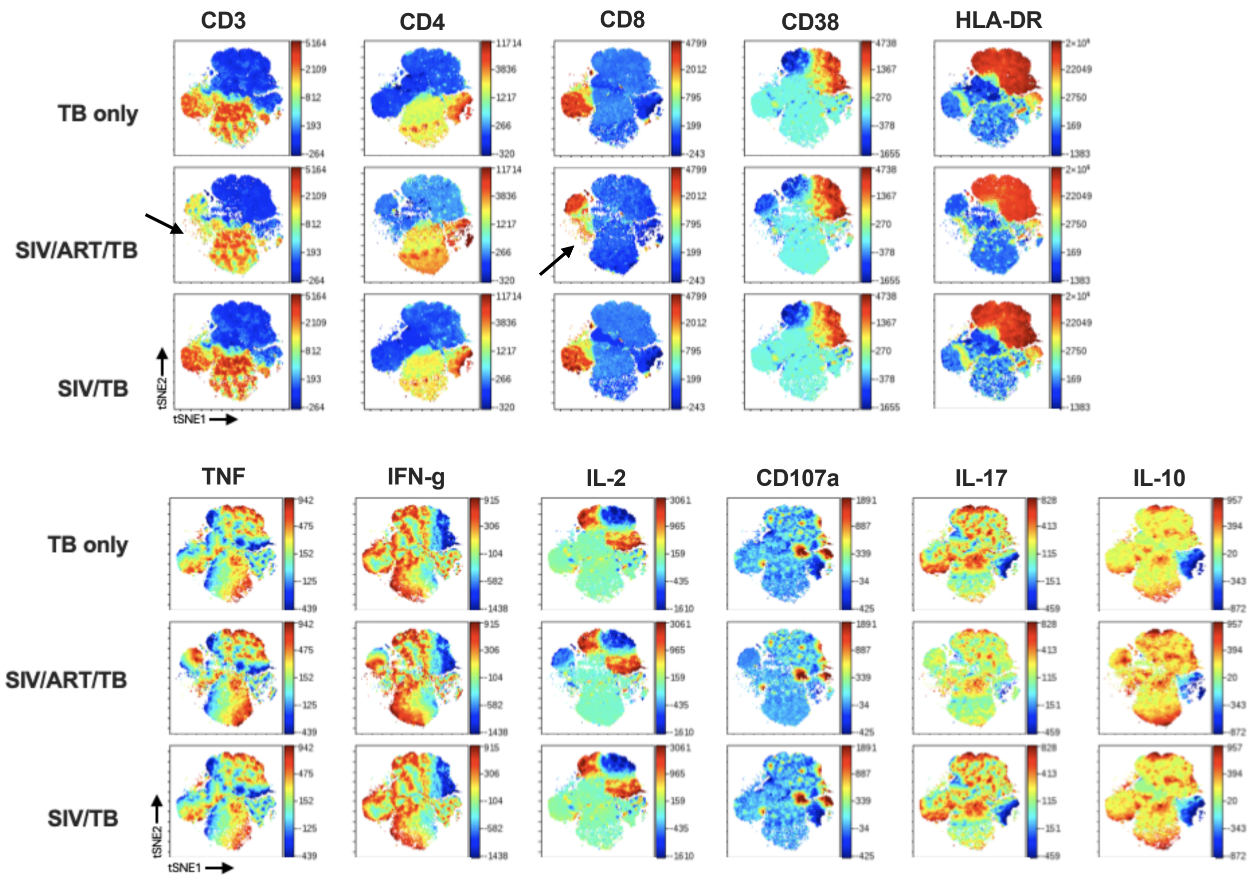


**Supplemental Figure 10**. t-SNE visualization of cellular markers lymphocytes in thoracic LNs. Pseudocolored maps show the relative expression of respective markers where red represents high expression and blue is low expression. Arrows highlight T cell populations based on phenotypic characteristics that differ between experimental groups. (n=12, 14, and 4 thoracic LN from 3 SIV/TB, 3 SIV/ART/TB, and 2 TB-only NHPs, respectively).

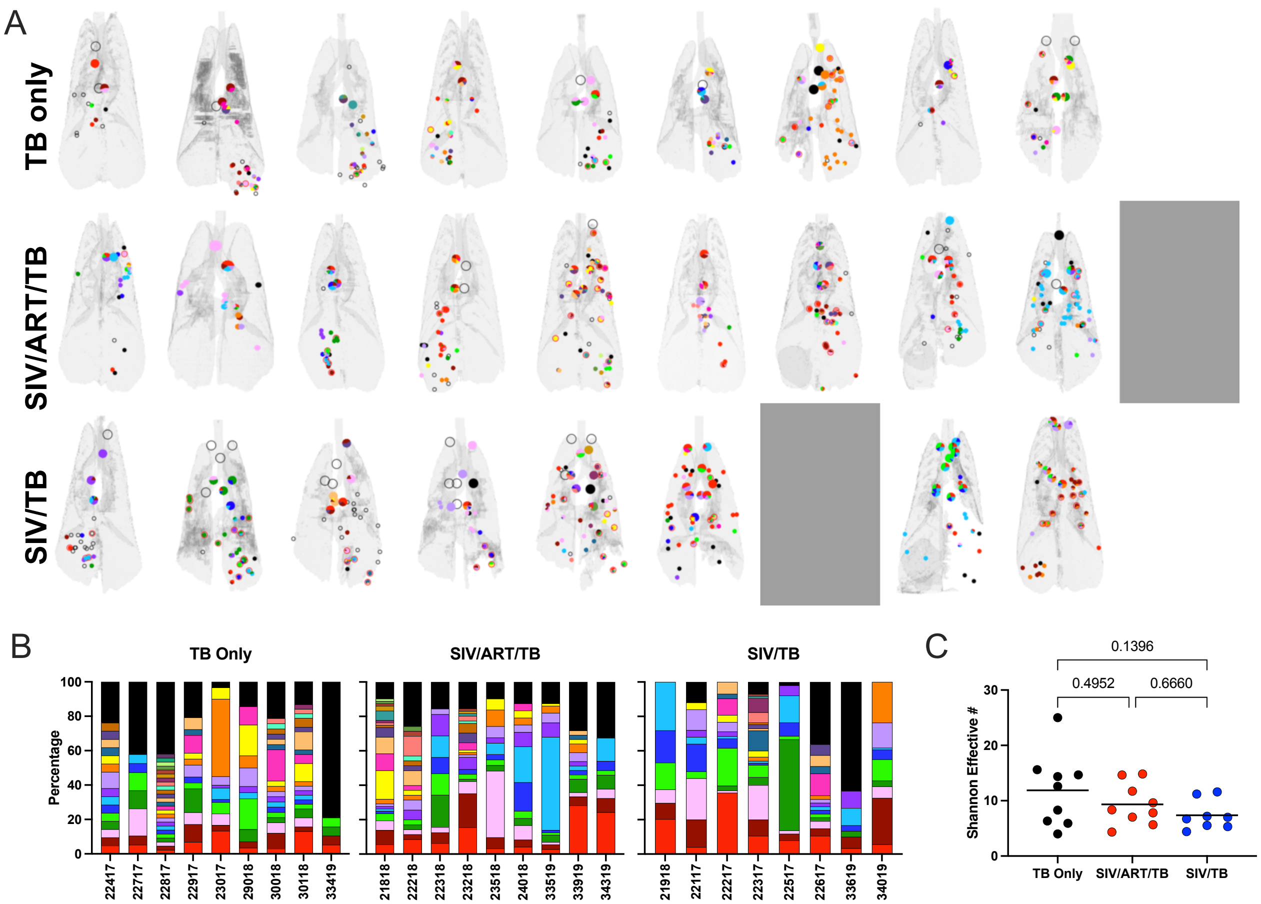


**Supplemental Figure 11**: A) Three-dimensional renderings of lungs with location of barcodes. Small dots represent granulomas; large circles represent thoracic lymph nodes; open circles represent sterile tissue. Two animals did not have viable barcodes; these are represented by gray boxes. B) Distribution of barcodes in the lungs each animal. Each color represents a barcode; black bars represent sterile samples. C) Diversity of barcodes (measured by the effective number quantified using the Shannon entropy index) in the lungs. Each dot represents an animal and lines represent group means. Groups were compared using ordinary one-way ANOVA with Tukey's multiple comparison adjusted p-values shown. TB only (n=9), SIV/ART/TB (n=9), SIV/TB (n=8).

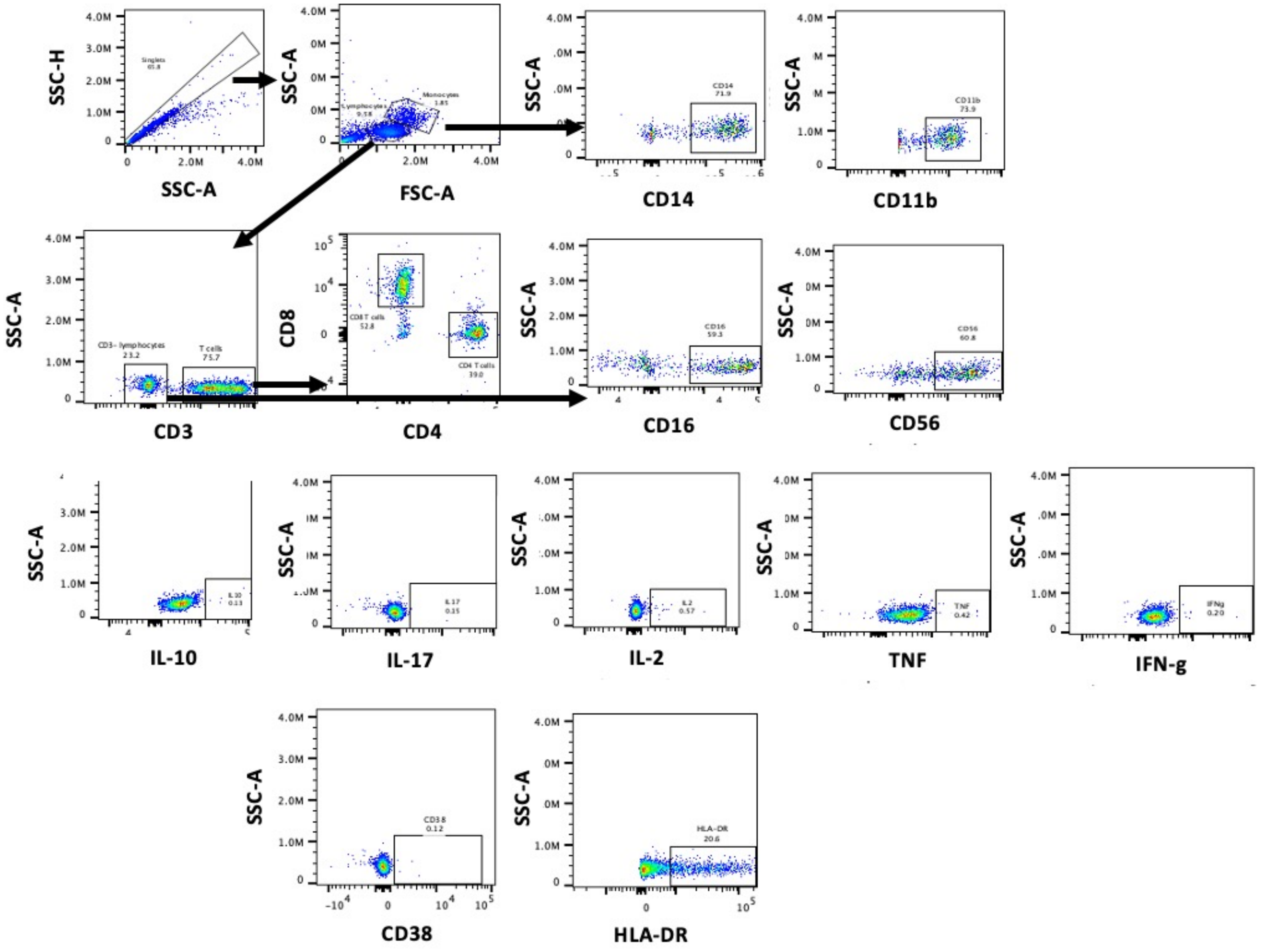


**Supplemental Figure 11.** Spectral flow gating strategy for PBMC and lymph node (CD107A) stimulated with ESAT6 and CFP10.

| Animal ID | Cohort | Gender | Age (yrs) | Weeks infected with SIV | Necropsy (wks post Mtb) |
|---|---|---|---|---|---|
| 22917 | TB-only | F | 8 | NA | 8 |
| 29018 | TB-only | M | 5 | NA | 12 |
| 30018 | TB-only | M | 5 | NA | 12 |
| 30118 | TB-only | M | 5 | NA | 12 |
| 33419 | TB-only | M | 6 | NA | 12 |
| 23017 | TB-only | F | 6 | NA | 13 |
| 22417 | TB-only | M | 6 | NA | 13 |
| 22717 | TB-only | M | 7 | NA | 13 |
| 22817 | TB-only | M | 7 | NA | 13 |
| 34419 | SIV/ART/TB | M | 7 | 26 | 10 |
| 34319 | SIV/ART/TB | M | 6 | 26 | 10 |
| 23218 | SIV/ART/TB | M | 8 | 27 | 11 |
| 33519 | SIV/ART/TB | F | 5 | 27 | 11 |
| 24018 | SIV/ART/TB | M | 9 | 27 | 11 |
| 23518 | SIV/ART/TB | M | 6 | 28 | 12 |
| 21818 | SIV/ART/TB | M | 5 | 28 | 12 |
| 22218 | SIV/ART/TB | M | 7 | 28 | 12 |
| 22318 | SIV/ART/TB | M | 6 | 29 | 13 |
| 33919 | SIV/ART/TB | M | 5 | 29 | 13 |
| 22217 | SIV/TB | M | 7 | 23 | 7 |
| 21918 | SIV/TB | M | 5 | 23 | 7 |
| 34019 | SIV/TB | M | 6 | 24 | 8 |
| 33719 | SIV/TB | M | 6 | 25 | 9 |
| 22317 | SIV/TB | M | 6 | 25 | 9 |
| 22117 | SIV/TB | M | 8 | 28 | 12 |
| 22517 | SIV/TB | M | 8 | 28 | 12 |
| 33619 | SIV/TB | F | 6 | 28 | 12 |
| 22617 | SIV/TB | M | 8 | 29 | 13 |
| 33819 | SIV-only | M | 6 | 11 | NA |
| 34119 | SIV-only | M | 6 | 29 | NA |
| 34219 | SIV-only | M | 6 | 29 | NA |

**Supplemental Table 1**. Animal identification number, experimental group, gender, age, weeks of SIV infection, and time to necropsy are shown.

### Supplemental Table 2

| Marker | Color | Clone | Company | Samples |
|---|---|---|---|---|
| CD11b | APC-Cy7 | ICRF44 | BD Bio Sciences | BAL |
| CD16 | PE-Cy5 | 3G8 | BD Pharmingen | BAL |
| CD206 | PerCPCy5.5 | 19.2 | BD Bio Sciences | BAL |
| CD3 | PE-CF594 | SP34 | BD Pharmingen | BAL |
| CD4 | BV510 | L200 | BD Horizons | BAL |
| CD45 | APC | D058 1283 | BD Bio Sciences | BAL |
| CD56 | PE-Cy5 | HCD56 | Biolegend | BAL |
| CD8 | Biotin:Streptavidin-AF700 | SK1 | BD Bio Sciences | BAL |
| HLA-DR | AF700 | G46 G | Fisher Scientific | BAL |
| IFN-g | FITC | B27 | BD Bio Sciences | BAL |
| IL-10 | efluor450 | JES34A3 | eBiosciences | BAL |
| TNF | FITC | Mab11 | BD Bio Sciences | BAL |
| CD11b | PECy7 | ICRF44 | BD Bio Sciences | PBMC |
| CD14 | BV768 | M5E2 | BD Bio Sciences | PBMC |
| CD16 | BUV563 | 3G8 | Biolegend | PBMC |
| CD3 | APC-Cy7 | SP34 | BD Pharmingen | PBMC |
| CD38 | PE | HIT2 | Biolegend | PBMC |
| CD4 | BV510 | L200 | BD Horizons | PBMC |
| CD56 | PeCy5 | FAB1059B | RD Systems | PBMC |
| CD8 | Biotin:Streptavidin-AF700 | SK1 | BD Bio Sciences | PBMC |
| HLA-DR | PE-CF594 | G46 G | Fisher Scientific | PBMC |
| IFN-g | AF700 | B27 | BD Bio Sciences | PBMC |
| IL-10 | efluor450 | JES34A3 | eBiosciences | PBMC |
| IL-17 | FITC | eBio64CAP17 | eBiosciences | PBMC |
| IL-2 | PerCPCy5.5 | MQ1-17H12 | BD Bio Sciences | PBMC |
| TNF | APC | Mab11 | BD Bio Sciences | PBMC |
| CD107a | PE | eBioH43 | H4A3 | Tissue |
| CD11b | PeCy7 | ICRF44 | BD Bio Sciences | Tissue |
| CD3 | APC-Cy7 | SP34 | BD Pharmingen | Tissue |
| CD38 | PeCy5 | HIT2 | Biolegend | Tissue |
| CD4 | BV510 | L200 | BD Horizons | Tissue |
| CD8 | Biotin:Streptavidin-AF700 | SK1 | BD Bio Sciences | Tissue |
| HLA-DR | PE-CF594 | G46 G | Fisher Scientific | Tissue |
| IFN-g | AF700 | B27 | BD Bio Sciences | Tissue |
| IL-10 | efluor450 | JES34A3 | eBiosciences | Tissue |
| IL-17 | FITC | eBio64CAP17 | eBiosciences | Tissue |
| IL-2 | PerCPCy5.5 | MQ1-17H12 | BD Bio Sciences | Tissue |
| TNF | APC | Mab11 | BD Bio Sciences | Tissue |

**Supplemental Table 2**. Spectral flow markers with respective fluorophore, clone, company and sample type shown.
